# Warming and resource enrichment decouple growth from enzymatic investment, shifting the competitive balance between native and invasive plants

**DOI:** 10.64898/2026.03.25.714348

**Authors:** Keren Yanuka-Golub, Roaa Abu-Alhof, Sawsan Hless, Jackline Abu-Nassar, Maor Matzrafi

## Abstract

Invasive plants can reshape ecosystems by altering soil biogeochemistry and microbial functioning under global change. Competitive interactions between the invasive *Conyza bonariensis* and the native *Helminthotheca echioides* were evaluated under warming, nitrogen enrichment, and elevated CO₂, together with rhizosphere microbial function in solitary versus competitive growth. Plants were grown alone or in interspecific competition under elevated temperature (27 vs 29 °C), ammonium–nitrate fertilization versus no fertilization, and ambient versus elevated CO₂ (400 vs 720 ppm). Plant traits and relative growth rate (RGR) were measured alongside potential extracellular enzyme activities (EEA) of α-D-glucosidase (C acquisition) and N-acetyl-β-D-glucosaminidase (NAGase; N acquisition) and functional gene abundances (nirS and bacterial amoA). To relate enzyme signals to plant demand and microbial biomass, we calculated a growth-normalized rhizosphere investment metric (Specific Rhizosphere Index; SRI) and a biomass-normalized investment metric (Specific Enzyme Activity; SEA). Competition effects were summarized as ΔSRI and ΔTax (change from alone to competition) to quantify how competition altered growth- and biomass-normalized investment. Plant responses were driver- and context-dependent. Elevated CO₂ produced the largest changes in growth traits, especially for the invasive species. Warming effects were modest in solitary plants but became apparent under competition, where elevated temperature reduced competitive suppression via increased invasive leaf production and reduced constraints on native leaf expansion. Fertilization caused comparatively small shifts in plant endpoints. Microbial responses depended strongly on soil conditioning history. Potential EEA showed limited shifts with warming and fertilization, whereas elevated CO₂ enhanced NAGase mainly in invasive-conditioned soils and increased nirS across soils. Despite overlap in ecoenzymatic stoichiometry, SRI and ΔTax revealed treatment- and legacy-dependent patterns in how competition re-scaled microbial C and N acquisition relative to plant growth and microbial biomass. Together, these results indicate that global change can decouple plant growth from enzymatic investment and reconfigure invasive–native interactions through shifts in above–belowground coupling.

## 1. Introduction

Biological invasions are a major driver of ecological change, with invasive plant species often differing fundamentally from native vegetation in their physiological, chemical, and functional traits (Drenovsky et al., 2012; Funk et al., 2016; Leishman et al., 2007; Te Beest et al., 2015). These differences can alter ecosystem processes, particularly those occurring in the soil (Belnap et al., 2005; Ehrenfeld, 2010, 2003). In contrast to many native species, invasive plants typically exhibit rapid growth rates, high specific leaf area, and elevated nutrient concentrations in their tissues. Such traits accelerate organic matter decomposition and intensify nitrogen cycling in the soil (Li et al., 2024). In addition, many invasive species exude distinctive secondary metabolites, including organic acids, antimicrobial compounds, and allelopathic chemicals, that restructure soil microbial communities, suppress native plant diversity, and confer competitive advantages to the invader (Kaštovská et al., 2015). Through these mechanisms, invasive plants modify soil chemistry, nutrient turnover, and plant-microbiome interactions, often disrupting ecological relationships essential for the persistence of native species (Weidenhamer and Callaway, 2010).

Environmental change, particularly global warming and rising CO_2_ levels, is expected to differentially affect invasive and native species (Ziska et al., 2019). Invasive plants often possess greater phenotypic plasticity, broader ecological tolerance, and faster growth rates, enabling them to exploit shifting climatic conditions more effectively than native species. Studies suggest that invaders may also benefit indirectly from water conserved by native plants during climate stress, thereby gaining an additional functional advantage (Blumenthal et al., 2013). Native species, often adapted to narrower ecological niches, may therefore decline in fitness under rapid environmental change. Such contrasts highlight the potential for climate warming to amplify invasion dynamics and undermine ecosystem resilience and biodiversity (Liu et al., 2017).

Nutrient enrichment, especially through nitrogen fertilization, is another global driver that disproportionately benefits invasive species. Nitrogen, a limiting nutrient for both plants and soil microorganisms, governs microbial metabolism, organic matter decomposition, and competition for soil resources. Invasive plants frequently respond more strongly to nitrogen additions than native species, increasing their biomass and competitive capabilities (Broadbent et al., 2018; Guo et al., 2023; Ren et al., 2019; Zhang et al., 2022). Fertilizers such as ammonium nitrate (NH₄NO₃), which supply both ammonium (NH₄⁺) and nitrate (NO₃⁻), accelerate carbon and nitrogen mineralization, stimulate soil respiration, and enhance greenhouse gas emissions, particularly CO₂ (Choi et al., 2011) and N_2_O (Qiu, 2015; Yu et al., 2021). These processes can further reinforce the success of nutrient-responsive invasive species (Bezabih Beyene et al., 2022; Lee et al., 2012; Weltzin et al., 2003; Yanuka-Golub et al., 2025). Invasive plants influence not only plant community structure but also elemental cycling in ecosystems. They typically exhibit higher nitrogen-use efficiency (NUE), elevated nitrogen concentrations in plant tissues, and lower carbon-to-nitrogen (C:N) ratios in their litter compared with native species (Jo et al., 2017; Sardans et al., 2017). These traits lead to faster decomposition rates, enhanced mineralization and nitrification, and greater availability of inorganic nitrogen (NH₄⁺, NO₃⁻) in the soil (Incerti et al., 2018).

Soil microbiomes play a central role in shaping the competitive balance between invasive and native plant species (Fahey and Flory, 2022). Invasive plants frequently alter soil properties such as pH, carbon content, and nutrient pools, which in turn restructure microbial community composition in ways that reinforce their own dominance (Torres et al., 2021). These shifts often create positive plant-soil feedbacks that enhance invader performance while reducing the growth of native species, as demonstrated for species including *Prosopis* and *Spartina* (Gao et al., 2022; Kaushik et al., 2023; Zhang et al., 2024). Environmental stressors such as drought further modulate these interactions, altering microbial community composition and the nature of plant–microbe feedbacks (Fahey and Flory, 2022; Wei et al., 2017). These findings support the view that soil microbial restructuring is not simply a consequence of invasion but a mechanism that actively promotes invasive species persistence and ecosystem-level change (Yanuka-Golub et al., 2025).

Soil enzymatic activity provides an important functional indicator of microbial responses to invasion. Invasive plants often increase the activity of enzymes involved in carbon, nitrogen, and phosphorus cycling, including, phenol oxidase, α/β-glucosidase, and N-acetyl-glucosaminidase (NAGase) (Aragón et al., 2014; Chen et al., 2025; Negesse et al., 2025; Zhou and Staver, 2019). NAGase, which hydrolyzes chitin into amino sugars, is closely associated with nitrogen mineralization and is commonly used as an indicator of fungal-mediated organic N turnover in soil (Ekenler and Tabatabai, 2004; Hoppe, 2018). In parallel, the abundance of key genes involved in nitrogen and phosphorus cycling, such as amoA (ammonia oxidation), *nirS* and *nirK* (denitrification), *nifH* (biological nitrogen fixation), and *phoD* (phosphorus mineralization), provides complementary insights into the functional potential of microbial communities under contrasting plant assemblies and environmental conditions (Bannert et al., 2011; Enebe and Babalola, 2021; Li et al., 2025). Here, gene abundance was evaluated as an independent indicator of microbial functional potential and was not integrated with enzyme activity in the index described below.

However, while absolute soil enzymatic activity and gene abundances provide a snapshot of microbial functional potential, they do not account for how strongly rhizosphere microbial activity is coupled to plant performance. To address this, we adopted an economic perspective on plant–microbe interactions (Bergmann et al., 2020; Wright et al., 2004) by calculating a Specific Rhizosphere Index (SRI) for two key enzymes, α-glucosidase and NAGase, as the ratio of potential enzyme activity to plant relative growth rate (RGR). In this framework, SRI reflects the magnitude of microbial investment per unit plant growth, with higher values indicating greater enzymatic activity relative to plant performance. To further assess the cost of plant–plant interactions (Emmett et al., 2020; Wen et al., 2022), we defined ΔSRI as a ’competition tax’ (Δ SRI): the additional metabolic cost a plant must incur to sustain growth when challenged by its competitive neighbor (Markham and Chanway, 1996). Positive ΔSRI values indicate an increased rhizosphere investment required to sustain growth under competition, representing a potential competition tax, whereas values near zero or below zero indicate little or no additional rhizosphere metabolic burden. Together, these metrics allow direct comparison of the competitive strategies of native and invasive species under shifting environmental conditions.

This methodological shift is particularly relevant given ongoing debates over the reliability of ecoenzymatic stoichiometry, which is typically inferred from a relatively small set of measurable enzymes and remains incompletely validated as a proxy for microbial nutrient limitation (Cui et al., 2022; Mori, 2024; Zheng et al., 2022). Identifying carbon and nutrient limitations in soil microorganisms remains a major challenge; however, by reconciling ecoenzyme and gene-based approaches with host growth trajectories, we can move toward a more mechanistic and quantitative understanding of plant-microbial nutrient limitation. This study aims to provide such insights, exploring how invasive plant-microbe interactions trigger adaptive mechanisms at the molecular and functional levels, ultimately uncovering the consequences of plant invasion for soil biogeochemical cycling.

Israel provides a unique context in which both native and invasive species co-occur across a range of habitats. A prominent invasive example is *Conyza bonariensis* (flaxleaf fleabane), an annual herb from the *Asteraceae* family, originating from tropical South America. Since its first documentation in the region in 1896, *C. bonariensis* has become one of the most widespread invasive plants in Israel (Dafni and Heller, 1982). Characterized by rapid growth, small wind-dispersed seeds, and prolific reproduction, it competes strongly with native species for soil moisture and nutrients, leading to reduced ecosystem stability and crop productivity. This species has also developed resistance to several key herbicides, making it highly difficult to control (Matzrafi et al., 2015). In contrast, *Helminthotheca echioides* (prickly ox-tongue) is a native annual species from the *Asteraceae* family. Covered with bristly hairs and flowering from May to October, it is common in moist habitats throughout northern and central Israel. The coexistence of native and invasive species such as these provides an opportunity to examine how environmental conditions influence competitive dynamics and belowground interactions.

We hypothesized that environmental stressors, specifically warming, elevated CO_2_, and nitrogen fertilization, would fundamentally alter the microbial economic efficiency of the rhizosphere. We predicted that the invasive species would maintain a ’Low-Cost, High-Output’ strategy (stable SRI), whereas the native species would exhibit a significantly higher ’competition tax’ (ΔSRI) under ambient conditions, a burden that may be alleviated by climatic shifts. To test this, we combined phenotypic measurements, extracellular enzyme assays, and qPCR targeting functional marker genes to link plant-driven differences in microbial dynamics with shifts in the surrounding soil environment.

## 2. Materials and Methods

### Soil properties

Bulk soil for the conditioning experiment was collected in the Newe Ya’ar region (northern Israel). Particle-size analysis showed 14.0% sand, 36.4% silt, and 49.6% clay. Prior to use, soil was air-dried, passed through a 2-mm sieve to remove coarse debris, and homogenized. This Mediterranean alluvial soil typically exhibits high water-holding capacity and slow drainage, factors likely to influence microbial activity and plant–soil interactions during the experiment.

### Seed pretreatment prior to sowing

Mature seeds of the native *H. echioides* and the invasive *C. bonariensis* were collected at the ridges of the Nahalal stream (32.707110, 35.185445). Seeds of each species were sown into sieved-homogenized Newe Ya’ar soil and maintained in a controlled-environment chamber at 20/27 °C (night/day), with a 14h photoperiod and 700µmol m^−2^ s^−1^ photosynthetic photon flux density (fluorescent lighting). Plants were grown under controlled conditions in a growth chamber (Conviron®, Gene 2000). Pots were watered regularly to maintain consistent soil moisture, avoiding both overly dry and waterlogged conditions. After germination, seedlings were grown until they developed at least four true leaves. Then, plants were moved to 1L pots filled with Newe Ya’ar soil. All treatments were applied one week after transplanting to make sure that plants had established properly.

### Experimental design of temperature, CO₂, and nitrogen fertilization experiments

To evaluate how changing environmental conditions modulate competitive interactions between the invasive plant *C. bonariensis* and the native species *H. echioides*, we conducted three controlled experiments manipulating temperature, atmospheric CO₂ concentration, and ammonium-nitrate fertilization. Temperature treatments consistent of 20/27 °C vs. 22/29 °C, and CO₂ levels were set at ambient (∼400 ppm) vs. elevated (∼720 ppm). To investigate the effect of agricultural run-off containing high concentrations of fertilization (N) on native-invasive competition soil feedbacks, fertilization was applied by adding ammonium nitrate to reach a final concentration of 25 mg N kg⁻¹ soil, while control plants received no nutrient additions. Each experiment included four planting treatments: native planted alone, invasive planted alone, native-invasive planted in the same pot, and soil-only as a control, established under identical growth conditions except for the imposed environmental factor. For all experiments, we monitored plant performance (leaf traits) weekly. The harvested soil from each pot was subsequently used for downstream microbial analyses. Several soil related measurements were also recorded: extracellular enzyme activities (α-glucosidase and N-acetyl-β-D-glucosaminidase), functional microbial genes via qPCR, and soil chemistry properties to assess nutrient availability and biochemical shifts. This integrated design allowed us to compare how each environmental driver differentially influences invasive-native plant competition and the functional responses of their associated soil microbial communities, relative to non-competitive conditions. Overall, the experiments combined two species combinations, three environmental conditions, and four biological replications per plant (five for the warming experiment), including control pots (soil only). In total, each growth chamber contained 16 pots (20 for the warming experiment). It should be noted that the experiments are not directly comparable due to their varying durations: the temperature and fertilization experiment ran for 35 days, and the CO₂ experiment for 28 days.

### Soil sampling and bulk soil chemical characterization

At the conclusion of each experiment, plants were manually removed from the pots and aboveground biomass was separated for phenotypic measurements. To access the root-associated soil while avoiding contamination from the green surface crust (algal/bryophyte layer) that had formed during the experiment, each pot was inverted to expose the root system. Based on the planting treatments, four distinct soil types were obtained: **Control soil**: Bulk soil from pots maintained without plants; **Native rhizosphere**: Interface soil conditioned solely by *H. echioides*; **Invasive rhizosphere**: Interface soil conditioned solely by *C. bonariensis*; **Competition rhizosphere**: A shared, integrated interface soil conditioned by both species. Due to the physical intertwining of root systems in the mixed-planting treatment, this fraction was collected as a single homogenized sample representing the combined influence of both the native and invasive plants.

For each type, soil adhering to and immediately surrounding the roots (or the equivalent depth in control pots) was gently dislodged and homogenized to obtain a bulk-rhizosphere interface subsample. Within each treatment group, these interface soils were processed to form representative samples for downstream microbial analyses, including DNA extraction and extracellular enzyme assays. The remaining soil in each pot, which was not in direct contact with the root system, was collected separately as bulk soil for moisture determination and chemical characterization. To determine soil moisture content, approximately 5 g of bulk soil were weighed and dried at 105 °C for 24 h. The resulting moisture values were used to normalize all enzymatic activity rates, gene abundance measurements, and chemical nutrient content to a dry-weight basis. Chemical analysis of the bulk soil included pH and inorganic nitrogen species (N-NH_4_ and N-NO_3_). pH was determined in a saturated soil paste extract following Standard Method SM 4500 H-B. Extractable nitrate N-NO_3_ was quantified using an aqueous extraction (1:5 w/v soil-to-deionized water) according to the American Society of Agronomy protocols (ASA, 1965; Method #1, Ch. 84-5). Ammonium N-NH_4_ was extracted using a 2N KCl solution (1:5 w/v) to displace exchangeable ions (Bremner, 1965), followed by colorimetric determination via the indophenol blue method (Kalra and Maynard, 1991). To ensure that nitrogen availability was not confounded by variations in soil moisture across environmental treatments (warming, CO_2_, and Nitrogen fertilization), all liquid extract concentrations (mg L^-1^) were normalized to a dry-soil mass basis (mg N kg^-1^ dry soil). This normalization accounted for the 1:5 extraction ratio and the gravimetric moisture content determined for each sample after drying at 105 °C for 24 h.

### DNA extraction and microbial gene abundance via qPCR

Whole-community genomic DNA was extracted from ∼480mg of homogenized root-associated fresh soil using the FastDNA™ SPIN Kit for Soil (MP Biomedicals) following the manufacturer’s protocol with minor modifications. DNA extracts were stored at −20 °C until further analysis. DNA purity and concentration were assessed using a NanoDrop ND-1000 spectrophotometer (Thermo Scientific, Wilmington, DE, USA). All PCR products were verified by 1.5% agarose gel electrophoresis, stained with Hy-View Nucleic Acid Stain (Cat. No. IMGS7011).

Absolute abundances of the bacterial 16S rRNA gene (proxy for total bacterial biomass) and selected functional genes were quantified by quantitative PCR (qPCR) following established protocols (Borchardt et al., 2021). Reactions were performed with SsoAdvanced™ Universal Inhibitor-Tolerant SYBR® Green Supermix (Bio-Rad) on a CFX96 Real-Time PCR System, and Cq values were processed in CFX Manager v2.3. Each 25 µL reaction contained 12.5 µL SYBR® Green Supermix, 0.3 µM of each primer, and 1 µL of DNA template. Because microbial biomass varied substantially among treatments, extracts were not normalized to a uniform DNA concentration; instead, a fixed template volume (1 µL) was used for all reactions. All samples and standards were run in technical triplicates. No-template controls (DNA-free water) were included on every plate. Assay specificity was confirmed during optimization by agarose gel electrophoresis (expected amplicon size) and monitored for every run by melt-curve analysis (65–95 °C) to ensure absence of primer dimers and non-specific amplification. Primer sequences, cycling conditions, and assay-specific amplification efficiencies are provided in Supplementary Table S1.

Several functional genes were initially screened by endpoint PCR (amoA, phoD, nirK, nirS, pmoA). Only nirS and amoA were consistently detected across all soil samples and were therefore quantified by qPCR. Standard curves and conversion of Cq to gene copies: For amoA and nirS, standard curves were generated using ten-fold serial dilutions (10⁻² to 10⁻⁹) of synthetic double-stranded DNA fragments (gBlocks; Integrated DNA Technologies) corresponding to the expected amplicon region (Han et al., 2023). gBlocks were suspended in TE buffer (stock 20 ng µL⁻¹). For each assay, Cq values of the standards were regressed against log_10_ of the standard concentration (in the same concentration unit used to prepare the dilution series; e.g., ng µL⁻¹). The resulting linear regression parameters (slope *m* and intercept *b*) were used to convert sample Cq to an equivalent DNA concentration (Equation 1): 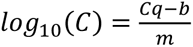

Where C is the DNA concentration of the target fragment in the extract (e.g., ng µL⁻¹). DNA mass was then converted to absolute copy number using the fragment length (bp):

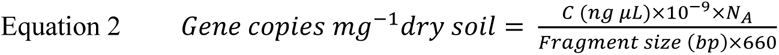

where N_A_ is Avogadro’s number (6.022×10²³ molecules mol⁻¹) and 660 g mol⁻¹ bp⁻¹ is the average molecular weight of one base pair of double-stranded DNA.

For the 16S rRNA gene absolute quantification, standard curves were generated from tenfold serial dilutions of purified genomic DNA of *Geobacter metallireducens* (**DSM 7210**), with the starting copy number calculated based on the precise mass (103 ng μl^-1^) of the purchased DNA and its known genome size (chromosome 3,997,420 bp + plasmid 13,762 bp ≈ 4.01 Mbp total; Aklujkar et al., 2009) and 16S copy number (n=2 obtained from rrnDB record): 16*S copies *μ*L*^−1^ = *genome copies μL*^−1^ × *n*.

For all primer sets, gene copy numbers were calculated for each technical replicate and then averaged (mean ± SD). Copy numbers were normalized to dry soil mass using the measured dry mass corresponding to each extraction and the total DNA elution volume (90 µL). The qPCR conditions and amplification efficiencies are described in Supplementary Table S1.

### Extracellular Enzyme Activity Assays

The activities of two common hydrolytic enzymes were measured to evaluate microbial potential for soil carbohydrate and nitrogenous organic-matter degradation using fluorometric assays. We quantified α-1,4-glucosidase (AG; EC 3.2.1.20), which hydrolyzes terminal α-glucosidic linkages in starch-like substrates, and β-1,4-N-acetylglucosaminidase (NAGase; EC 3.2.1.52), which cleaves N-acetyl-β-D-glucosamine from chitin and peptidoglycan. Assays followed modified protocols from German et al. (2011), using Sigma-Aldrich substrates 4-methylumbelliferyl α-D-glucopyranoside (69591) for AG and 4-methylumbelliferyl N-acetyl-β-D-glucosaminide (M2133) for NAGase. Three technical replicates of each sample were assayed per substrate concentration, along with negative controls (no sample). Enzyme activities were quantified by the release of 4-Methylumbelliferone (MUF) from fluorogenic substrates. Standard curves were prepared fresh daily using 4-Methylumbelliferone sodium salt (MW = 198.15 g/mol), dissolved in sterile deionized water to a stock concentration of 600 µM, and serially diluted to generate a standard range of 100–600 µM. For the purpose of final activity calculations, standard concentrations were corrected to reflect the molar mass of neutral MUF (MW = 176.17 g/mol), as this represents the actual fluorophore released during enzymatic hydrolysis.

Soil slurries (homogenates) were prepared fresh on the day of analysis by vortexing 2.0 g (± 0.2) of fresh soil in 10 mL of sterile deionized water at half speed for 10 minutes (Vortex Genie 2, Scientific Industries) to obtain a homogeneous suspension. Several control treatments were included: (1) homogenate control (soil and water without substrate), (2) substrate control (substrate and water without soil), (3) homogenate quality control (homogenate spiked with a known MUF concentration), (4) standard MUF control (MUF in water), and (5) blank control (water only). All reactions (samples, controls, and standards) were conducted in a total volume of 2 mL, composed of 1 mL soil homogenate/water and 1 mL substrate solution, and incubated in the dark at room temperature for 1 hour. After incubation, tubes were centrifuged at 14,000 rpm for 5 minutes. A volume of 250 µL of the supernatant was transferred to a black 96-well microplate, and 50 µL of sterile water was added to bring the total volume to 300 µL.

Fluorescence was measured using a microplate reader (Feyond-A300) with excitation at 365 nm and emission at 450–460 nm. Fluorescence values were converted to product concentrations using the corrected standard curve, with quenching corrections applied based on individual sample quench curves. Final enzyme activity was expressed as µmol MUF released per gram of dry soil per hour (µmol g⁻¹ h⁻¹), accounting for total reaction volume, incubation time, dry soil weight equivalent, and background fluorescence (German et al., 2011). Detailed standard curve parameters and quenching values are provided in Supplementary Table S2.

### Integrated assessment of plant performance and aboveground-belowground feedbacks

To synchronize aboveground physiological performance with belowground functional shifts, several proxies provided an estimate of physiological performance, capturing the developmental trajectories of both species across two distinct biotic contexts: individuals grown in a solitary state and those subjected to interspecific competition (Markham and Chanway, 1996). The growth of both native *H. echioides* and invasive *C. bonariensis* was quantified via a rosette development index, following established vitality metrics for this species (Karkanis et al., 2022): G = Leaf Number (N) * Rosette Diameter (D). These temporal dynamics were integrated into a unified Relative Growth Rate (RGR), calculated as:

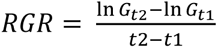, where t_1_ and t_2_ are the initial and final experimental time points, respectively.

To evaluate microbial nutrient-mobilization relative to plant performance, we calculated a Specific Rhizosphere Index (SRI) for two key enzymes: α-glucosidase and β-1,4-N-acetylglucosaminidase (NAGase). The SRI was defined as the ratio of potential enzyme activity to plant relative growth rate (RGR), representing the microbial investment in rhizosphere nutrient acquisition per unit plant growth. Higher SRI values therefore indicate a stronger soil enzymatic response relative to plant performance, whereas lower values indicate tighter coupling between plant growth and microbial activity (Dijkstra et al., 2013; Emmett et al., 2020; Huo et al., 2017; Meier et al., 2020). This integrated approach allowed for a direct assessment of whether environmental drivers induced a functional coupling or decoupling between plant nutrient demand and microbial supply. To quantify the metabolic burden on the microbial community, we calculated the Specific Investment Cost (Tax). Potential activities for α-glucosidase and NAGase were normalized to 16S rRNA gene copies, providing a per-capita estimate of microbial resource investment (Sinsabaugh et al., 2013). This ’tax’ was then compared across solitary and competitive contexts to determine the net shift (Δ_Tax_) in microbial effort induced by interspecific plant interactions.

## Results

### 3.1 Plant physiological responses under warming, fertilization, and elevated CO₂

Across the experiments, invasive and native plants differed in their growth responses, and these responses depended strongly on whether plants were grown alone or in direct competition. In the warming experiment, temperature effects were small under ambient conditions and when plants were grown alone, but became more pronounced under competition, where warming relieved competitive stress and narrowed the difference between solitary and competitive plants (Fig. 1, Table S3).

**Figure 1.**
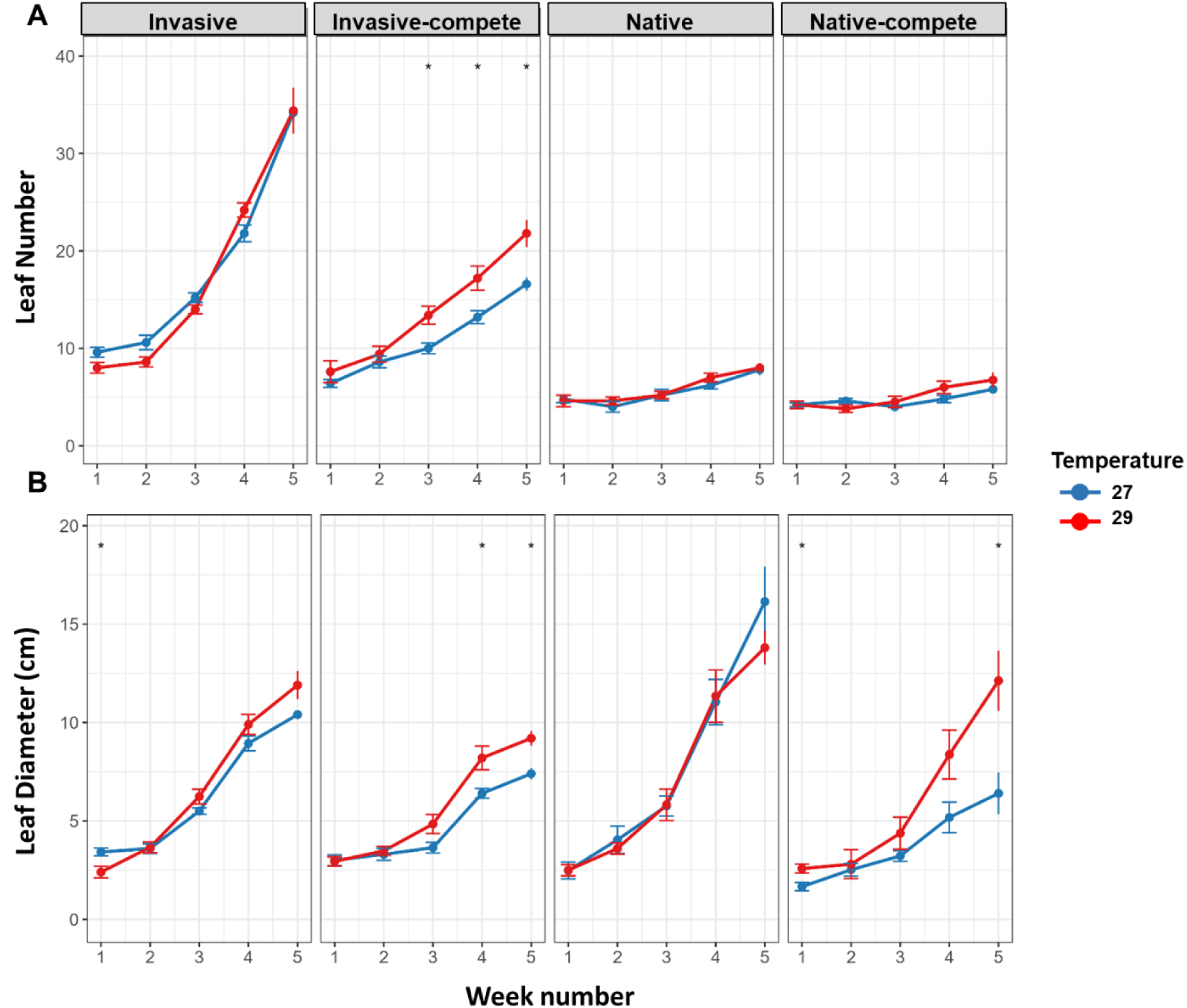
Temporal dynamics of plant growth under temperature and competition treatments. Mean (± SE) leaf number (A), and leaf diameter (B) were monitored weekly over 35 days for invasive and native plants grown alone or under competition (Invasive_Compete, Native_Compete) at 27 °C (blue) and 29 °C (red). Statistical differences between temperatures within each plant type and time point are indicated above the curves analyzed using t-test (p < 0.05).

For leaf production (Fig. 1A), invasive *C. bonariensis* grown alone produced similarly high leaf numbers at both temperatures by week 5 (34.2 ± 1.6 at 27 °C vs. 34.4 ± 5.2 at 29 °C), indicating limited direct warming effects under non-competitive conditions. Under competition, however, warming increased invasive leaf number (21.8 ± 3.1 at 29 °C vs. 16.6 ± 1.5 at 27 °C, week 5), thereby reducing the magnitude of the competitive penalty at elevated temperature. Conversely, *H. echioides* showed minimal temperature dependence in leaf production, regardless of whether plants were grown alone or in competition (alone: 7.8 ± 1.3 vs. 8.0 ± 1.0; competition: 5.8 ± 0.45 vs. 6.75 ± 1.71, week 5).

Leaf diameter (Fig. 1B) clearly showed that warming reduced competitive constraints on growth, especially for the native species. In *C. bonariensis*, warming modestly increased final leaf diameter both when plants were grown alone (11.9 ± 1.6 at 29 °C vs. 10.4 ± 0.4 at 27 °C) and under competition (9.2 ± 0.8 vs. 7.4 ± 0.7), also reducing the gap between solitary and competitive plants at the higher temperature. In *H. echioides*, competition strongly constrained leaf expansion at 27 °C (6.4 ± 2.4 under competition vs. 16.1 ± 4.0 alone), but this constraint was largely alleviated at 29 °C, where competitive plants reached leaf diameters much closer to those of solitary plants by week 5 (12.1 ± 3.4 under competition vs. 13.8 ± 1.9 alone).

Plant growth responses to CO₂ enrichment depended strongly on species identity and competitive context (Fig. 3; Table S3). When grown alone, *C. bonariensis* produced substantially more leaves than the native species (Fig. 3A), and elevated CO₂ was associated with a marked increase in final leaf number relative to ambient CO₂ (34.75 ± 6.75 vs. 20.25 ± 2.63 leaves). Under competition, this positive CO₂ effect on invasive leaf number persisted but was reduced in magnitude (19.25 ± 3.95 vs. 14.75 ± 0.96 leaves), indicating that biotic interactions constrained the CO₂ stimulation of invasive leaf production.

Compared with the invasive species, *H. echioides* exhibited only minor CO₂-driven changes that differed between solitary and competitive pots. When grown alone, the native plant showed only a small increase in leaf number under elevated CO₂ (9.25 ± 0.50 vs. 8.75 ± 0.96 leaves). Under competition, however, leaf number was lower under elevated CO₂ than under ambient CO₂ (7.00 ± 1.15 vs. 8.25 ± 1.26 leaves), suggesting that the direction of the CO₂ effect on native leaf production depended on competitive context. Notably, this contrasts with the warming experiment, where native leaf number remained broadly similar across temperatures in both solitary and competitive pots, implying that leaf production in the native species is relatively temperature-insensitive but can shift modestly under CO₂ enrichment depending on competition.

Leaf diameter responses further highlighted species-specific strategies (Fig. 3B). In *C. bonariensis*, differences in leaf diameter between CO₂ treatments were modest when plants were grown alone (11.28 ± 0.94 at elevated CO₂ vs. 13.77 ± 8.00 at ambient CO₂) and slightly higher under elevated CO₂ when grown in competition (7.28 ± 1.38 vs. 5.83 ± 0.43). In contrast, *H. echioides* exhibited a strong CO₂-linked increase in leaf expansion when grown alone, with substantially larger leaves under elevated CO₂ (37.23 ± 4.24 vs. 26.98 ± 3.42). This enhancement was not observed under competition, where leaf diameters were comparable or slightly lower under elevated CO₂ (23.05 ± 3.45 vs. 25.20 ± 1.62). Together, these results indicate that CO₂ enrichment primarily promoted invasive leaf production, whereas the native response was expressed mainly through enhanced leaf expansion in solitary plants, with competition dampening these CO₂-driven growth responses (Fig. 3; Table S3).

In the fertilization experiment, ammonium–nitrate addition (25 mg N kg⁻¹ soil) produced only modest shifts in plant growth, and treatment differences remained small at week 5 (Fig. 2; Table S3). For the invasive *C. bonariensis*, competition consistently reduced growth relative to plants grown alone, lowering both leaf number and leaf diameter under both nutrient regimes (Fig. 2A–B). Fertilization partially mitigated competitive suppression of *C. bonariensis*: under competition, fertilized plants ended with more leaves than unfertilized plants (21.0 ± 4.0 vs. 17.0 ± 3.2) and slightly larger leaves (7.0 ± 0.6 vs. 5.88 ± 1.03), although these trends were not significant by week 5 (Table S3).

**Figure 2.**
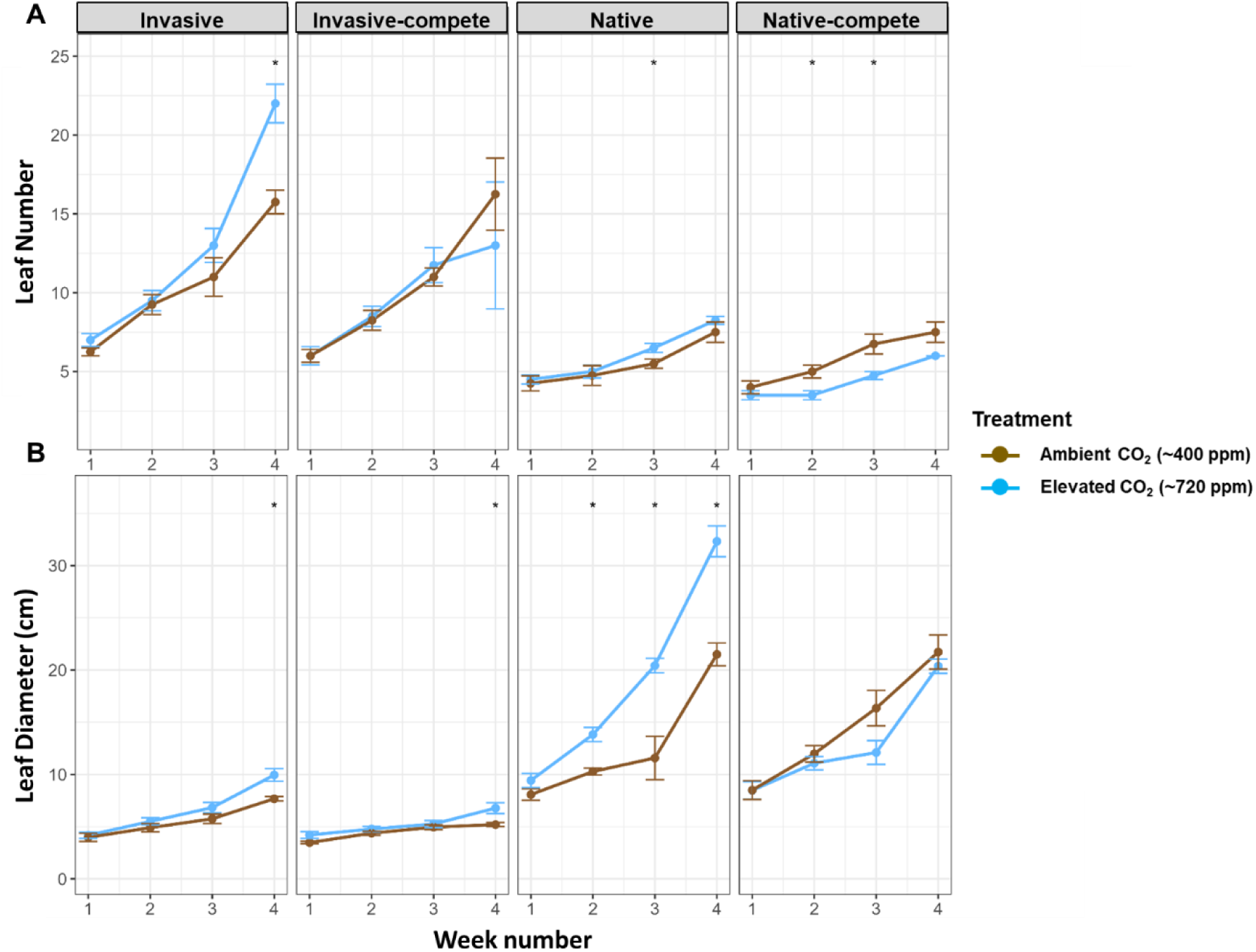
Temporal dynamics of plant growth responses to elevated CO₂ and competition. Mean (± SE) leaf number (A) and leaf diameter (B) were monitored weekly over 28 days for invasive and native plants grown alone or under interspecific competition. Plants were exposed to ambient (Low CO₂, Brown) or elevated CO₂ (High CO₂, Blue) conditions. Statistical differences between temperatures within each plant type and time point are indicated above the curves analyzed using t-test (p < 0.05).

**Figure 3.**
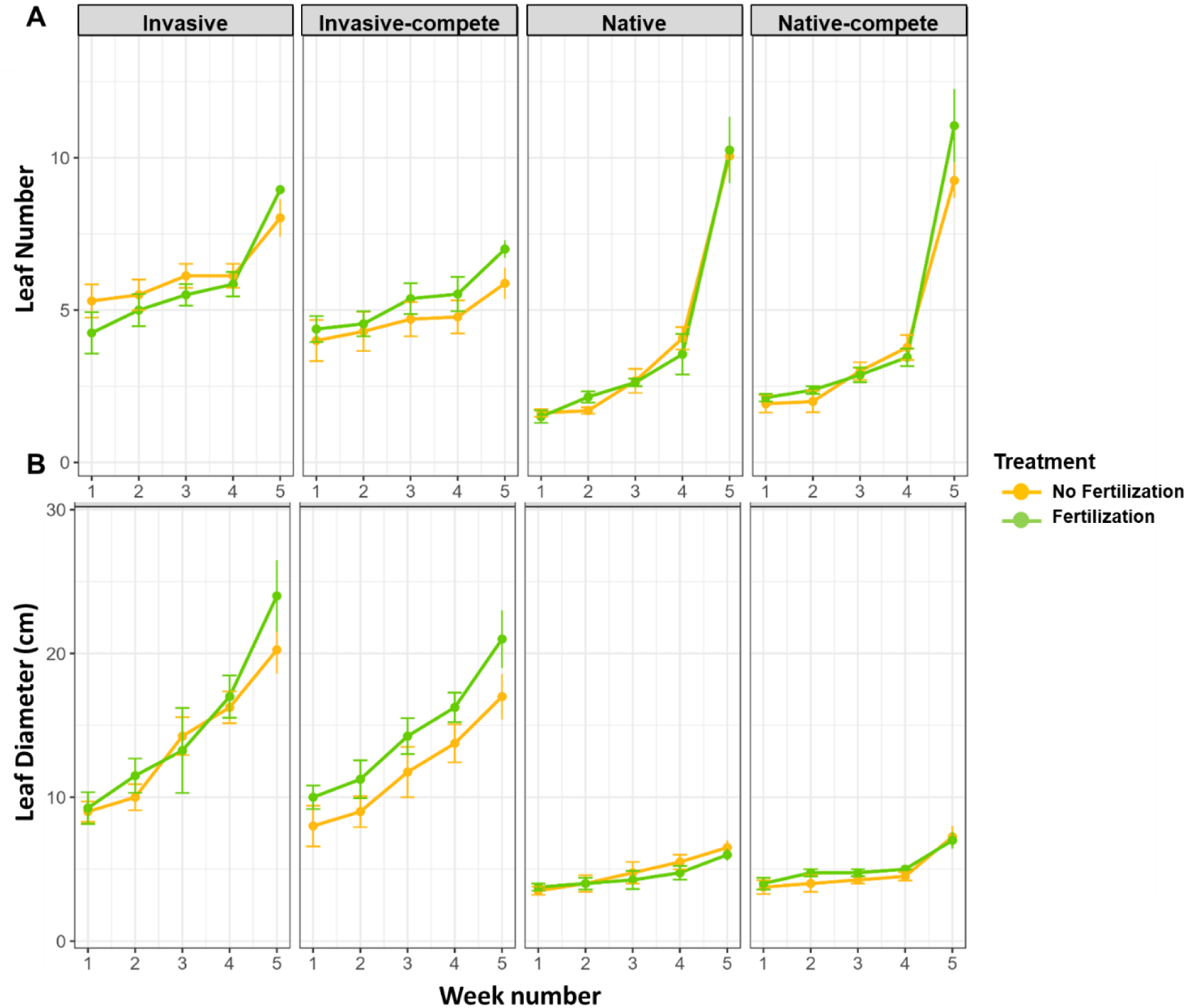
Temporal dynamics of plant growth under fertilization and competition. Mean (± SE) leaf number (A) and leaf diameter (B) were monitored weekly over 35 days for invasive and native plants grown alone or under competitive conditions (Invasive_Compete, Native_Compete). Plants were grown with fertilization (green) or without fertilization (orange). Statistical differences between temperatures within each plant type and time point are indicated above the curves analyzed using t-test (p < 0.05).

By contrast, *H. echioides* showed minimal fertilization effects on leaf number, and competitive plants had a slightly higher leaf number than solitary plants under both nutrient regimes (fertilized: 7.00 ± 1.15 vs. 6.00 ± 0.82; unfertilized: 7.25 ± 1.50 vs. 6.50 ± 1.00; Fig. 2A; Table S3). Native leaf diameter responses were small and variable, with only minor differences between fertilized and unfertilized treatments and between solitary and competitive growth (Fig. 2B).

Trait-specific responses, however, revealed contrasting sensitivities between species. In the native *H. echioides*, fertilization was associated with a comparatively steep increase in leaf number during the later stages of the experiment, indicating enhanced leaf production rather than increased leaf expansion (Fig. 2A). In contrast, fertilization effects in *C. bonariensis* were more strongly expressed through increases in leaf diameter, particularly in plants grown alone, suggesting preferential allocation toward leaf expansion rather than leaf initiation (Fig. 2B). Despite these divergent trait responses, nutrient addition had little effect on native growth under competitive conditions, where leaf number and leaf diameter trajectories remained nearly identical between fertilized and unfertilized treatments. Overall, fertilization did not override competitive hierarchies, but instead revealed species-specific growth strategies: native plants responded primarily via increased leaf number, whereas invasive plants exhibited greater plasticity in leaf expansion.

### 3.2 Soil physicochemical conditions across experimental treatments

Baseline soil physicochemical conditions varied across plant-soil legacies and experimental manipulations (Table 1). Raw data, ART ANOVA outputs, and letter groupings derived from Holm-adjusted art.con pairwise contrasts (α = 0.05) are provided in Supplementary Table 3.

**Table 1.**
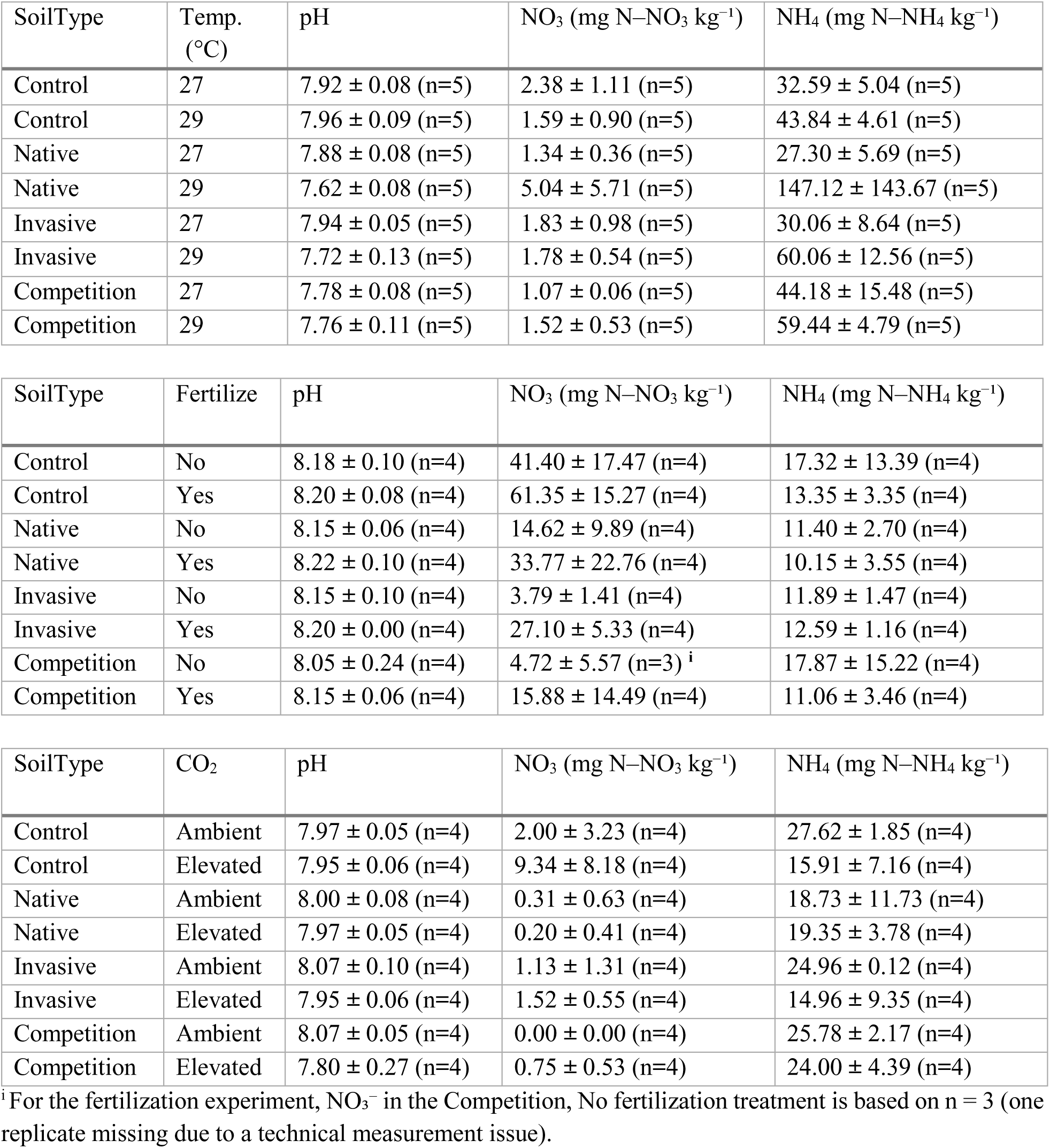
Soil chemical characteristics across plant–soil treatments. Values are mean ± SD.

#### Temperature experiment

Across soil legacies, warming from 27 to 29 °C was generally associated with higher NH₄⁺ (mg N–NH₄ kg⁻¹ dry soil), with the strongest shift in native-conditioned soils (27.30 ± 5.69 to 147.12 ± 143.67) and more moderate increases in the control treatment with no plants grown (32.59 ± 5.04 to 43.84 ± 4.61), invasive (30.06 ± 8.64 to 69.69 ± 24.14), and competition soils (44.18 ± 15.48 to 59.44 ± 4.79; Table 1). NO₃⁻ (mg N–NO₃ kg⁻¹ dry soil) remained low across most soil types and temperatures (≈1.07–2.38), but was higher and markedly more variable in native soils at 29 °C (5.04 ± 5.71; Table 1). pH showed modest but consistent legacy-dependent differences, with higher values in control and invasive soils at 27 °C (7.92 ± 0.08 and 7.94 ± 0.05) and lower values in invasive and native soils at 29 °C (7.72 ± 0.13 and 7.62 ± 0.08; Table 1).

#### Fertilization experiment

Soil pH was stable across legacies and between fertilized and unfertilized treatments (Table 1). NH₄⁺ also showed little separation among soil types or fertilization treatments. In contrast, NO₃⁻ clearly differentiated soils and responded to fertilization: control soils maintained the highest nitrate concentrations overall, and fertilized treatments showed higher nitrate than unfertilized treatments across soil legacies (Table 1; Supplementary Table 3).

#### CO₂ experiment

Elevated CO₂ was associated with a small reduction in pH, most apparent in competition soils (Ambient: 8.07 ± 0.05 vs Elevated: 7.80 ± 0.27; Table 1). NO₃⁻ patterns were strongly legacy-dependent and CO₂-sensitive: control soils showed higher nitrate under elevated CO₂ (Ambient: 2.00 ± 3.23 vs Elevated: 9.34 ± 8.18), whereas competition soils remained near zero under both CO₂ treatments (Table 1). NH₄⁺ was also lower under elevated CO₂, with the clearest contrast in control soils (Ambient: 27.62 ± 1.85 vs Elevated: 15.91 ± 7.16; Table 1). Compact letters summarizing significant pairwise differences are reported in Supplementary Table 3.

### 3.3 Ecoenzyme-based evidence for microbial C and N limitation under warming, fertilization, and elevated CO₂

Warming, fertilization, and elevated CO₂ differentially influenced potential extracellular enzyme activities (EEA) and functional gene abundances, with patterns strongly mediated by soil conditioning history (Supplementary Fig. S4A-B; Supplementary Table 4). Temperature emerged as the dominant driver across all soil types, consistent with the well-established temperature sensitivity of microbial N-acquisition processes (Allison and Treseder, 2008; Sinsabaugh et al., 2009). Potential EEA for both enzymes remained broadly similar between 27 and 29 °C across soil types (Supplementary Fig. S4A), while legacy effects were consistent: native- and invasive-conditioned soils generally exhibited higher potential activities than control and competition soils. The gene response to warming was more specific, where *nirS* increased in control soils at 29 °C, such that control soils largely closed the gap with the other legacies, whereas *nirS* in the remaining soil types stayed similar to their 27 °C levels (Supplementary Fig. S4B; Supplementary Table 4). The fertilization experiment produced no clear shifts in either potential EEA or the abundances of *nirS* and bacterial *amoA* across soils or treatments (Supplementary Fig. S4A-B; Supplementary Table 4). In contrast, elevated CO₂ effects were soil-type specific, enhancing β-1,4-N-acetylglucosaminidase activity primarily in invasive-conditioned soils, while having little to no effect in native, control, or competition-conditioned soils (Supplementary Fig. S4A). In parallel, *nirS* abundance increased under elevated CO₂ across soil types (with control soils relatively high under ambient CO₂), indicating a consistent CO₂-associated increase in denitrification potential (Supplementary Fig. S4B; Supplementary Table 4). This pattern suggests that plant legacies amplify microbial responsiveness to altered carbon inputs under CO₂ enrichment, likely through changes in substrate availability and microbial nutrient demand (Cheng et al., 2014; Phillips et al., 2012).

#### 3.3.1 Linking plant growth to rhizosphere enzyme investment

Because plant growth and rhizosphere microbial processes are tightly coupled, via root-derived carbon inputs, microbial enzyme-mediated nutrient mobilization, and feedbacks on plant nutrient supply, we linked aboveground growth rates to belowground function using two complementary, theory-grounded normalizations of enzyme activity (Bengtson et al., 2012; Dijkstra et al., 2013). First, we calculated a growth-normalized rhizosphere investment index (here termed the Specific Rhizosphere Index; SRI), defined as potential enzyme activity divided by plant relative growth rate (RGR). This formulation captures the enzyme investment per unit plant growth, aligning with rhizosphere theory that plant growth and nutrient demand can intensify belowground processes (focusing here on enzyme production) and thereby couple plant performance to microbial nutrient mobilization (Dijkstra et al., 2013; Kuzyakov and Razavi, 2019). Second, we quantified biomass-specific microbial investment (here termed Tax, computed as Specific Enzyme Activity; SEA), defined as potential enzyme activity normalized by microbial biomass, where biomass was estimated here using 16S rRNA gene copy number (Sinsabaugh et al., 2009).

#### 3.3.2 Warming reconfigures competition effects on growth- and biomass-normalized investment

Using the growth-normalized investment index (SRI) and biomass-normalized (16S abundance) microbial investment (SEA), Figure 4 shows that the impact of competition between the native and invasive plants on belowground C- and N-acquisition investment was temperature-dependent and differed between soil legacies. At 27 °C, competition was associated with a pronounced decrease in SRI in invasive-conditioned soils for both α-D-glucosidase (C acquisition; Fig. 4A) and NAGase (N acquisition; Fig. 4C), whereas native-conditioned soils showed only small shifts. In parallel, ΔTax decreased under competition in the invasive legacy (Fig. 4A, C), indicating reduced per-biomass enzyme investment (SEA/16S) from alone to competition. Together, these patterns indicate that under ambient temperature the invasive legacy exhibits a stronger competition-linked reorganization of enzyme investment relative to plant growth and microbial biomass for both C and N acquisition pathways.

**Figure 4.**
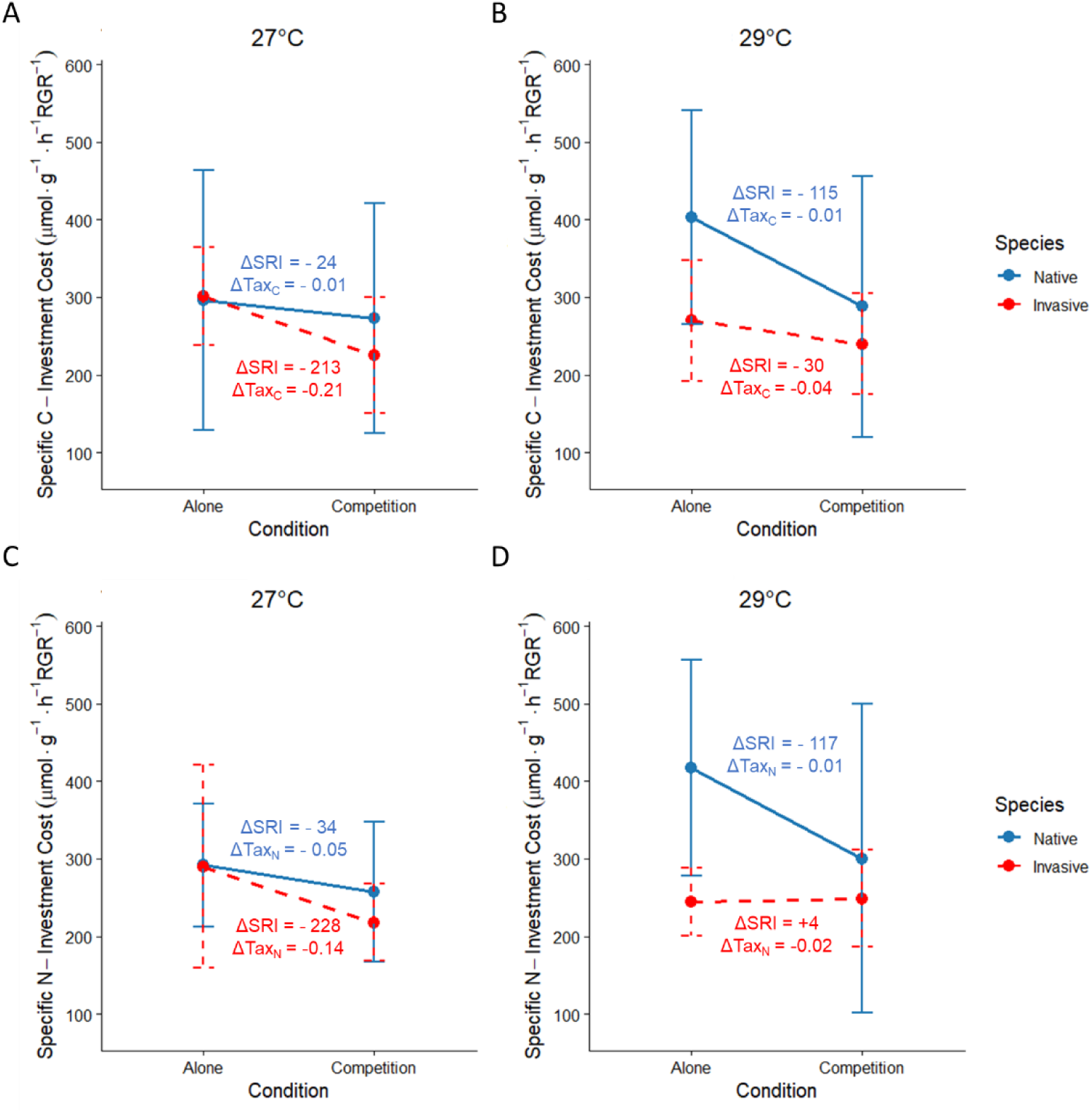
Competition effects on microbial enzyme-investment traits for Carbon based on α-D-glucosidase activity (A, B) and Nitrogen based on β-1,4-N-acetylglucosaminidase (NAGase) activity (C, D) acquisition across temperature treatments. The Specific Rhizosphere Index (SRI) values are plotted for Alone vs Competition conditions for Native (blue) and Invasive (red) treatments; points are treatment means connected within species, with error bars (95% CI). Text annotations quantify the within-species shift between solitary (Alone) and competitive conditions: ΔSRI, defined as the slope between the Alone and Competition means after converting enzyme activity to a growth-normalized investment cost (enzyme activity divided by relative growth rate, RGR), and ΔTax, defined as the change in biomass-normalized enzyme investment (Specific Enzyme Activity, SEA), calculated as enzyme activity normalized by microbial biomass (16S rRNA gene copy number).

At 29 °C, the competition response shifted toward native-conditioned soils (Fig. 4B, D). Native soils showed large competition-associated decreases in SRI for both enzymes, while invasive soils displayed only modest change in α-D-glucosidase-based SRI and little to no change in NAGase-based SRI. Notably, ΔTax values were near zero at 29 °C for both legacies, suggesting that the competition-associated changes in SRI under warming were not primarily driven by changes in per-biomass enzyme investment, but rather by changes in the balance between potential enzyme activity, microbial biomass, and plant growth across treatments. Supporting EEA and gene abundance patterns are provided in Supplementary Fig. S4A-B and Supplementary Table S4 and S5.

Ecoenzymatic stoichiometry provides a complementary view of microbial resource allocation by comparing relative investment in C- vs N-acquiring enzymes based on potential activities. In the ln-ln space of α-D-glucosidase (C acquisition) versus NAGase (N acquisition), points falling above the 1:1 line indicate proportionally greater investment in C acquisition (a “C-limited” domain), whereas points below the line indicate proportionally greater investment in N acquisition (an “N-limited” domain) (Eco-stoichiometry plot; Supplementary Table 4C). The enzyme C:N index (Supplementary Table 4C) captures this relationship numerically, where the 27 °C and 29 °C samples largely overlap in this enzyme-allocation space, consistent with the limited temperature effect on raw potential enzyme activities (Supplementary Fig. S4A). Thus, the strong temperature-dependent patterns in Figure 4 are best interpreted as changes in the coupling between belowground potential function and aboveground growth (SRI) and in biomass-normalized investment (SEA), rather than a wholesale shift in the relative allocation between C- and N-acquiring enzymes (Supplementary Fig. S4A–B; Supplementary Table 4C).

#### 3.3.3 CO₂ enrichment reconfigures competition effects on investment indices

Figure 5 shows that CO₂ enrichment reconfigured how competition translated into belowground C- and N-acquisition investment in the native vs invasive rhizospheres. Under ambient CO₂, competition increased both C-investment cost (α-D-glucosidase; Fig. 5A) and N-investment cost (NAGase; Fig. 5C), with the largest increase in the invasive rhizosphere (ΔSRI_*C*_ = +428; ΔSRI_*N*_ = +395), whereas the native rhizosphere showed smaller increases (ΔSRI_*C*_ =+77; ΔSRI_*N*_ = +75). These competition-driven increases in SRI under ambient CO₂ coincided with negative ΔTax values (SEA/16S), indicating reduced per-biomass enzyme investment from alone to competition growth mode despite higher growth-normalized costs (Fig. 5A, C; supporting EEA and gene patterns in Supplementary Fig. S4A–B and derived indices in Supplementary Table 4). Under elevated CO₂, competition effects were reversed (Fig. 5B, D). In the native rhizosphere, competition shifted toward lower investment costs (ΔSRI*C* = −39; ΔSRI_*N*_ = −40), while in the invasive rhizosphere the competition effect was modest for C acquisition (ΔSRI_*C*_ = +92) and reversed for N acquisition (ΔSRI_*N*_ = −59). Across panels, ΔTax values were small under elevated CO₂, suggesting that CO₂ enrichment reduced the extent to which competition-driven changes were accompanied by large shifts in biomass-normalized microbial enzyme investment (SEA/16S) (Fig. 5; Supplementary Table 4). Ecoenzymatic stoichiometry was evaluated using the same ln–ln framework described above (α-D-glucosidase vs. NAGase; 1:1 reference line). Under ambient and elevated CO₂, samples largely overlapped in this enzyme-allocation space, showing no clear CO₂-driven shift relative to the 1:1 line (Supplementary Table 4C). This indicates that CO₂ enrichment did not substantially re-partition potential enzyme investment between C- and N-acquisition pathways. Accordingly, the CO₂-dependent responses in Figure 5 are best interpreted as changes in the coupling between potential belowground function and aboveground growth (SRI) and in biomass-normalized investment (SEA/16S) across competitive contexts, rather than changes in enzyme allocation per se (Supplementary Fig. S4A–B; Supplementary Table 4).

**Figure 5.**
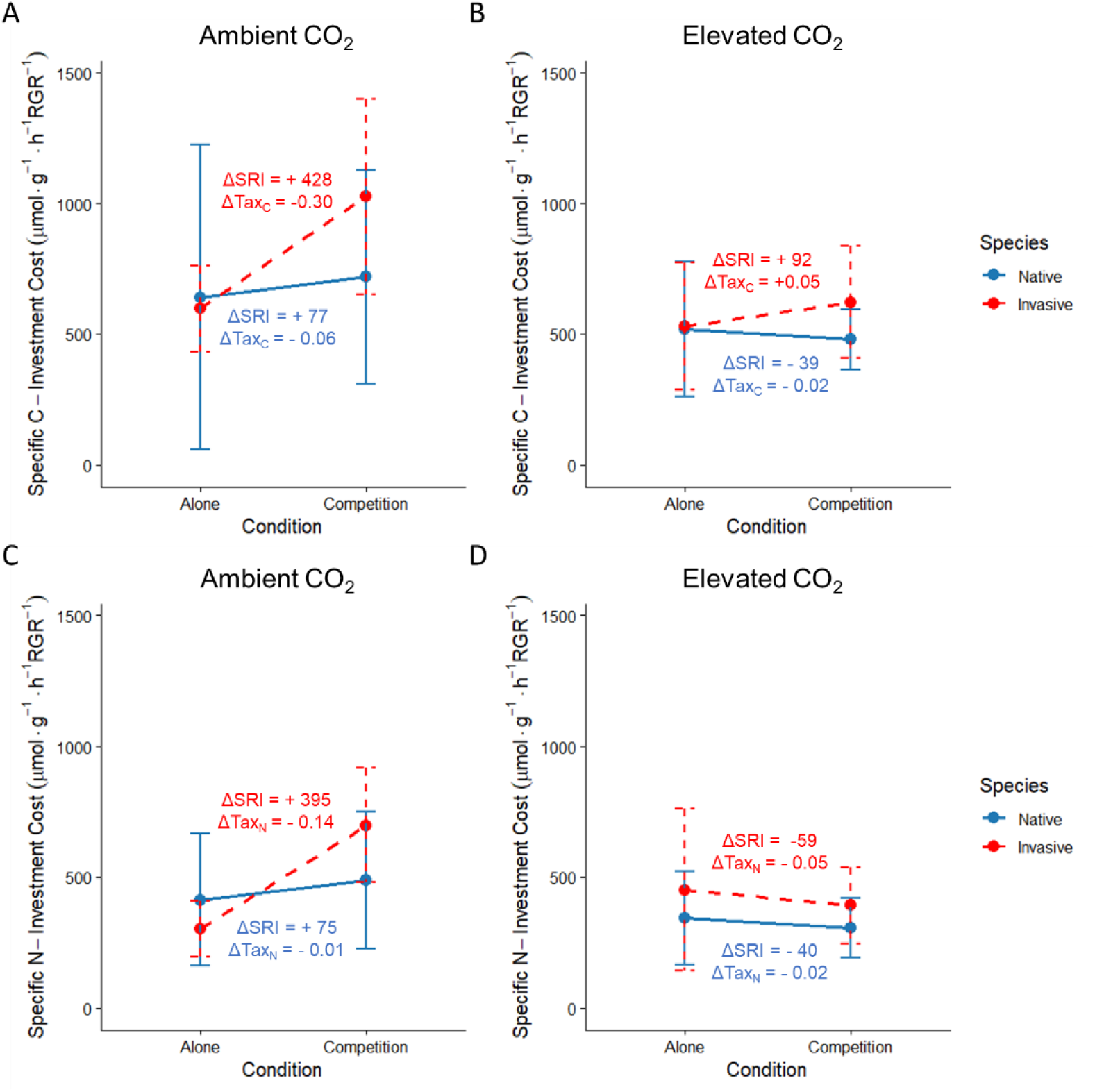
Competition effects on microbial enzyme-investment traits under CO₂ treatments. SRI is plotted for Alone vs Competition for Native (blue) and Invasive (red). SRI and ΔTax are shown for C acquisition (α-D-glucosidase; A, B) and N acquisition (NAGase; C, D) under ambient (Low CO₂) and elevated (High CO₂) conditions. Points are treatment means (±95% CI) for Alone vs. Competition within each soil legacy (native- vs invasive-conditioned), connected within legacy; text annotations report ΔSRI (slope between Alone and Competition means for EEA/RGR) and ΔTax (change in SEA/16S between Alone and Competition).

#### 3.3.4 Fertilization modestly reshapes competition-linked enzyme investment costs

Figure 6 shows that fertilization changed how competition translated into growth-normalized enzyme investment costs (SRI) and biomass-normalized investment (ΔTax; SEA/16S) for both C- and N-acquisition enzymes. Under no fertilization, competition generally reduced SRI in the invasive rhizosphere for both α-D-glucosidase (ΔSRI_C_ = − 620; Fig. 6A) and NAGase (ΔSRI_N_ = − 335; Fig. 6C), while effects in the native rhizosphere were smaller (ΔSRI_C_ = −137; ΔSRI_N_ = +38). In contrast, under fertilization, competition increased SRI for both species (Fig. 6B, D), with the strongest response for invasive C acquisition (ΔSRI_C_ = +896; ΔSRI_N_ = +338) and positive shifts also observed for the native rhizosphere (ΔSRI_C_ = +310; ΔSRI_N_ = +102).

**Figure 6.**
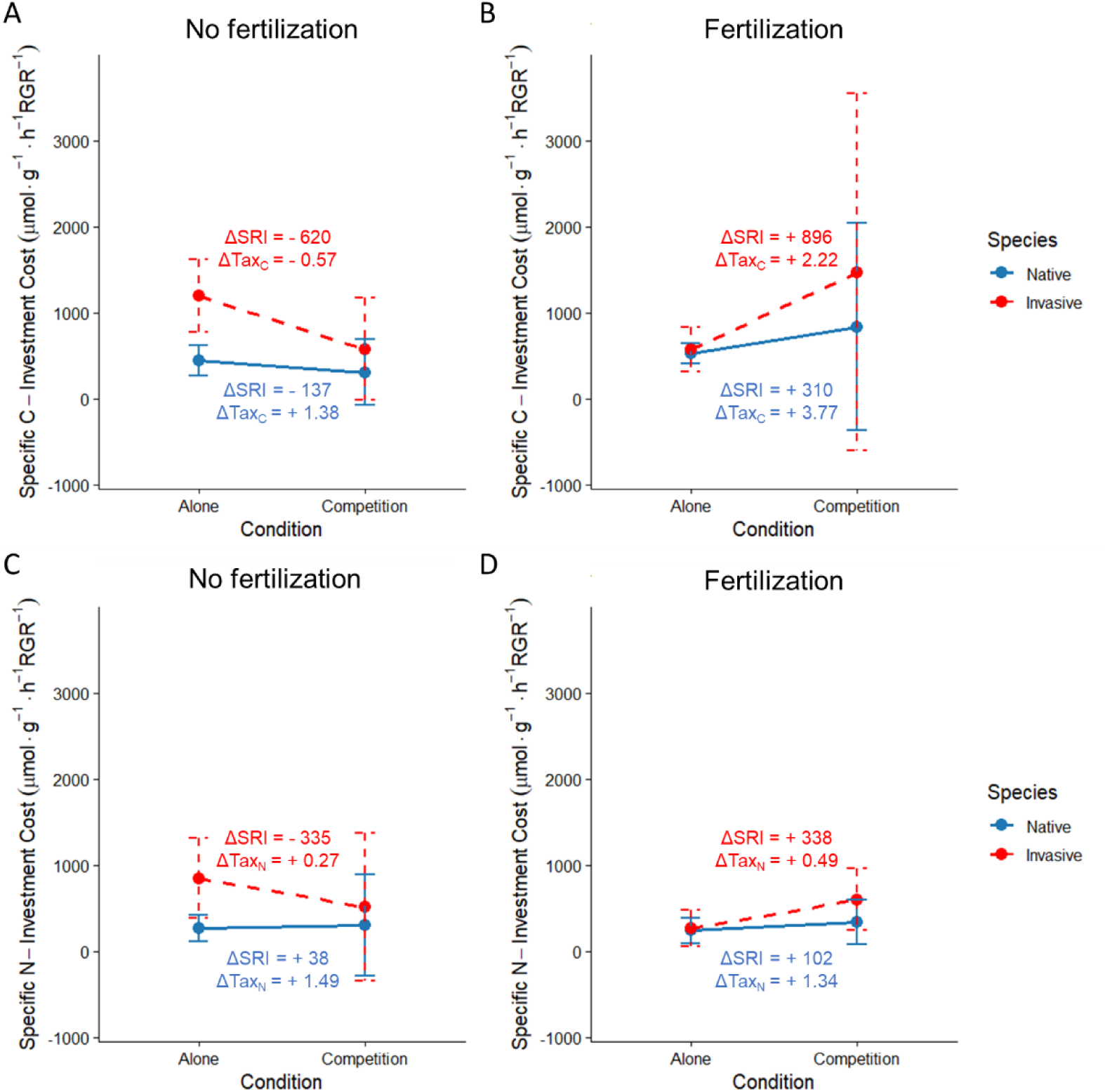
Competition effects on microbial enzyme-investment traits under fertilization treatments. SRI is plotted for Alone vs Competition for Native (blue) and Invasive (red). SRI and ΔTax are shown for C acquisition (α-D-glucosidase; A, B) and N acquisition (NAGase; C, D) under no fertilization and fertilization (NH₄NO₃ addition). Points are treatment means (±95% CI) for Alone vs Competition within each soil legacy (native- vs invasive-conditioned), connected within legacy; text annotations report ΔSRI (EEA/RGR slope) and ΔTax (SEA/16S change) from Alone to Competition.

Patterns in ΔTax indicated that these competition-driven shifts in growth-normalized costs were accompanied by different microbial investment responses between species. Across fertilization treatments, the native rhizosphere showed consistently positive ΔTax values (ΔTax_C_ = +1.38 to +3.77; ΔTax_N_ = +1.34 to +1.49), indicating increased per-biomass enzyme investment from solitary to competitive growth. In the invasive rhizosphere, ΔTax was near zero or negative under no fertilization (ΔTax_C_ = −0.57; ΔTax_N_ = −0.27) and remained small but positive under fertilization (ΔTax_C_ = +2.22; ΔTax_N_ +0.49). Together, Fig. 6 indicates that fertilization shifted competition effects toward higher growth-normalized enzyme investment costs, while native rhizospheres showed the clearest increase in biomass-normalized investment under competition. Ecoenzymatic stoichiometry (ln α-glucosidase vs. ln NAGase) clustered predominantly above the 1:1 line, consistent with relatively greater C-acquisition potential across samples (Supplementary Fig. S4C). Fertilized and unfertilized treatments overlapped extensively in this space, indicating no clear fertilization-driven shift in relative allocation between C- and N-acquiring enzymes.

Across treatments, plant traits indicate how warming, fertilization, and CO₂ alter competitive outcomes, whereas SRI and ΔTax show how rhizosphere enzyme investment is re-scaled relative to plant growth and microbial biomass. Notably, these indices showed clear treatment- and context-dependent patterns even when raw potential enzyme activities and final growth endpoints changed only modestly.

## Discussion

The accelerating pace of global environmental change is fundamentally restructuring plant communities by altering the balance of competitive interactions between native and invasive species. Invasive plants often possess a suite of functional traits, including high phenotypic plasticity, rapid resource acquisition, and superior biomass allocation, that allow them to capitalize on environmental perturbations more effectively than their native counterparts (Davidson et al., 2011; Pyšek et al., 2020). As atmospheric CO_2_ concentrations rise and global temperatures shift, these "opportunistic" traits may be further amplified, potentially expanding the niche breadth of invasive taxa and facilitating their dominance in novel climates (Hellmann et al., 2008; Sorte et al., 2013). However, the degree to which these species maintain their advantage depends heavily on the presence of other biotic factors, such as inter-species competition, which can either constrain or exacerbate the physiological benefits provided by a changing environment (Bradley et al., 2010; Ziska et al., 2019).

Our results demonstrate that environmental change modulates invasive-native interactions primarily through its effects on plant physiological performance, and that these shifts depend strongly on competitive context. Among the drivers tested, elevated CO_2_ elicited the most pronounced influence on growth traits, particularly for the invasive *C. bonariensis*. Under non-competitive conditions, elevated CO₂ was associated with a marked increase in invasive leaf number, whereas under competition this stimulatory effect persisted but was reduced, indicating that biotic interactions constrained the CO₂ benefit. In contrast, native responses to elevated CO₂ were comparatively modest and more context dependent: leaf number changed only slightly when plants were grown alone and tended to shift in the opposite direction under competition, suggesting that CO₂ enrichment alone is unlikely to overturn competitive hierarchies but can reinforce existing asymmetries when combined with biotic stress. This aligns with recent findings that elevated CO_2_ disproportionately increases the biomass of invasive species compared to natives (Bajwa et al., 2019; Sobuj et al., 2024). Notably, the native CO₂ response was expressed more clearly through leaf expansion in solitary plants, whereas invasive responses were expressed mainly through leaf production, consistent with the idea that invasive success can be reinforced via distinct trait pathways rather than uniform increases in all growth metrics.

Warming and fertilization produced more nuanced outcomes, but their effects were again strongly competition dependent. Under warming, temperature effects were small when plants were grown alone, yet under competition elevated temperature reduced competitive penalties, increasing invasive leaf production and relaxing constraints on native leaf expansion (Fig. 1). This pattern aligns with the broader view that warming can shift competitive balance by altering growth costs and resource capture under interspecific stress (Dukes and Mooney, 1999; Liu et al., 2017; Sorte et al., 2013). Fertilization, by comparison, generated only modest shifts in final growth endpoints, but it tended to buffer invasive performance under competition (e.g., slightly higher leaf number and diameter in competitive *C. bonariensis*), consistent with the fluctuating resource hypothesis that invaders can disproportionately exploit resource pulses (Davis et al., 2000; Jin et al., 2025). Importantly, fertilization did not produce a uniform invasive advantage; instead, it highlighted trait-specific sensitivities that differed from warming. Under nutrient enrichment, responses were expressed mainly through invasive leaf expansion and comparatively subtle shifts in native leaf number, whereas warming and CO₂ more consistently aligned with invasive leaf-number responses and native leaf-diameter responses under specific competitive contexts. Together, this reversal in trait sensitivity across drivers underscores that plant functional responses are driver-specific, rather than universally aligned along an invasive–native axis. Nonetheless, even these subtler responses tended to favor the invasive species, particularly under competition, consistent with evidence that invaders can disproportionately exploit resource enrichment and climatic warming (Dukes and Mooney, 1999; Sorte et al., 2013; Liu et al., 2017; Davis et al., 2000; Jin et al., 2025).

Notably, native responses to CO₂ were strongly contingent on competition: while isolated native plants produced slightly more leaves under elevated CO₂, native plants in competitive pots showed higher leaf numbers under ambient CO₂. This context-dependent reversal supports the view that CO₂ enrichment alone is unlikely to overturn competitive hierarchies, but instead interacts with biotic stress in ways that can reinforce existing asymmetries (Butler et al., 2025).

These plant-level shifts have direct implications for belowground processes, as changes in growth rate, allocation, and tissue traits are tightly coupled to rhizosphere carbon inputs and microbial functioning. Accordingly, we examine how these environmentally driven differences in plant performance translate into alterations in soil microbial activity and functional potential, providing a mechanistic link between aboveground dominance and ecosystem-level biogeochemical responses.

Mechanistically, our results suggest that invasive–native growth responses under global change are accompanied by shifts in the efficiency and strategy of rhizosphere nutrient mobilization, and that these shifts depend strongly on both abiotic context (warming vs. CO₂ enrichment vs. N addition) and biotic context (competition) (Dijkstra et al., 2013; Pugnaire et al., 2019; Van der Putten et al., 2013). Across experiments, potential enzyme activities and functional gene abundances (Supplementary Fig. S4A–B) reflect the baseline functional capacity of the soil microbiome. In contrast, the growth- and biomass-normalized investment metrics in Figs. 4–6 show how that capacity is expressed in relation to plant performance, by quantifying changes in coupling between aboveground growth and belowground nutrient-acquisition potential.

Integrating plant traits (Figs. 1–3) with these investment metrics (Figs. 4–6) reveals a consistent pattern: environmental drivers altered competitive outcomes through trait-specific shifts in growth (leaf production vs. leaf expansion), and these shifts were accompanied by driver-specific re-scaling of enzyme “costs” relative to plant demand (SRI = enzyme/RGR) and microbial biomass (ΔTax; SEA/16S).

Under warming, competitive penalties were reduced aboveground (Fig. 1). Belowground, responses were expressed mainly as temperature-dependent shifts in growth–enzyme coupling and in the soil legacy where competition most strongly altered investment costs (Fig. 4). Raw potential enzyme activities differed strongly by plant legacy and competition treatment, with higher activity in native- and invasive-conditioned soils than in unplanted controls and generally higher activity under solitary than competitive growth (Supp. S4A). Thus, warming did not simply increase enzyme activity. Instead, it changed how enzyme activity scaled with plant growth (SRI), consistent with studies showing that warming can alter competitive outcomes and that warming effects on soil enzymes depend on microbial biomass, enzymatic traits, and resource context rather than temperature alone (Lu et al., 2016; Fanin et al., 2022). Because enzyme activities reflect microbial metabolic demand, and specific activities can reveal changes that bulk measurements may obscure, the stronger warming signal in SRI points to a reorganization of growth–investment coupling rather than a simple increase in enzyme production (Caldwell, 2005; Fanin et al., 2022; Lu et al., 2016).

Under elevated CO₂, the clearest aboveground stimulation occurred in the invasive species, which produced more leaves when grown alone; under competition, this benefit persisted but was weaker (Fig. 3). Native responses were more context dependent, with CO₂ mainly promoting leaf expansion in solitary plants and competition reversing that response. Belowground, competition increased SRI costs under ambient CO₂, especially in the invasive rhizosphere, but these effects were attenuated under elevated CO₂ (Fig. 5). Because ΔTax values remained small and enzyme stoichiometry overlapped strongly (Supp. Figure 4C), the main CO₂ effect appeared to be a shift in how competition translated into growth-normalized microbial investment, rather than a major reallocation between C- and N-acquiring enzymes. This interpretation is consistent with studies showing that elevated CO₂ can enhance invasive plant performance more strongly in monoculture than in mixture, or leave competitive outcomes only weakly changed, while at the same time increasing belowground C inputs and root exudation that reshape soil/rhizosphere enzyme activity and nutrient cycling in a context-dependent manner (Dong et al., 2021; Hager et al., 2020; Kelley et al., 2011; Xu et al., 2013).

Fertilization produced comparatively modest changes in plant endpoints (Fig. 3), yet it altered the direction of competition effects on SRI and revealed contrasting per-biomass investment responses between species (Fig. 6), illustrating that coupling can shift even when plant traits change only subtly. Across drivers, ecoenzymatic stoichiometry showed substantial overlap among treatments (Supplementary Fig. S4C; Supplementary Table 4C), reinforcing the interpretation that the dominant signal was not a wholesale shift in C vs. N allocation, but rather changes in how enzyme potential is leveraged relative to growth and biomass. Together, these integrated patterns support a model in which global change modifies invasion outcomes by reshaping the efficiency of rhizosphere nutrient acquisition per unit plant performance, and by altering whether competition is expressed mainly through shifts in biomass-normalized microbial investment (SEA/16S) versus shifts in growth coupling (enzyme/RGR).

Finally, the gene-based patterns provide an important constraint on interpretation. Increases in *nirS* under elevated CO₂ (Supplementary Fig. S4B) suggest that CO₂ enrichment can increase denitrification potential across soil legacies, even when enzyme responses are legacy-specific. This decoupling between enzyme and gene responses underscores the need for integrative metrics such as SRI and SEA, which help distinguish changes in potential capacity (enzyme and gene pools) from changes in functional coupling between plants and microbes. In this context, control soils showing higher N-acquisition efficiency (Supplementary Fig. S6; Supplementary Table 4C) may reflect legacy-dependent differences in the balance between upstream depolymerization and downstream N-cycling potential, reinforcing the idea that invasion outcomes depend not only on plant traits, but also on how soil legacies modulate the microbial pathways that supply nutrients to plants under competition and global change.

Competition-conditioned soils generally exhibited reduced or intermediate enzyme responses across treatments, indicating constrained microbial nitrogen acquisition under combined biotic and abiotic stressors. In contrast, α-D-glucosidase activity was largely structured by soil type and showed weak or non-significant responses to temperature and CO₂, reflecting greater functional stability of carbon-acquiring enzymes relative to nitrogen-acquiring enzymes across environmental gradients (Sinsabaugh et al., 2009; German et al., 2012). Collectively, these results demonstrate that plant legacy effects modulate microbial functional sensitivity to global change drivers, with invasive conditioning selectively enhancing microbial responsiveness to CO₂ while warming exerts a more universal control across soil types. Such legacy-dependent responses highlight the potential for invasive plants to reshape belowground nutrient cycling under future climate scenarios (Strickland et al., 2013; Bardgett & Wardle, 2010).

## Data availability

Data will be made available upon reasonable request.

## Declaration of generative AI and AI-assisted technologies in the manuscript preparation process

During the preparation of this work, the authors used Gemini (Google) and ChatGPT (OpenAI) in order to assist with text revision, language polishing, and improving the organization and clarity of the manuscript. These technologies were specifically utilized to refine the conceptual framework of the "Microbial Tax" and to synthesize the relationship between potential enzyme activity and the Specific Rhizosphere Index (SRI). After using these tools, the authors reviewed and edited the content as needed and take full responsibility for the content of the published article.

## Acknowledgments

Authors would like to thank Aseel Sadeq for her valuable contribution. We gratefully acknowledge the Israel Ministry of Innovation, Science and Technology for funding project number 4673 (Investigating greenhouse gases emissions in plant invasion hotspots as a model for aquatic ecosystem restoration and management), which supported the research efforts leading to the development and writing of this paper.

## Competing interests

The authors declare no competing interest.

## Author contributions

K.Y.-G. and M.M. conceived and designed the study. R.A.-A., S.H., and J.A.-N. performed the experiments and collected the data. K.Y.-G., M.M., and J.A.-N. drafted and revised the manuscript.

## Notes

### Competing Interest Statement

The authors have declared no competing interest.

## References

Aklujkar, M., Krushkal, J., DiBartolo, G., Lapidus, A., Land, M.L., Lovley, D.R., 2009. The genome sequence of Geobacter metallireducens: features of metabolism, physiology and regulation common and dissimilar to Geobacter sulfurreducens. BMC Microbiol. 9, 109.

Allison, S.D., Treseder, K.K., 2008. Warming and drying suppress microbial activity and carbon cycling in boreal forest soils. Glob. Chang. Biol. 14, 2898–2909.

Aragón, R., Sardans, J., Peñuelas, J., 2014. Soil enzymes associated with carbon and nitrogen cycling in invaded and native secondary forests of northwestern Argentina. Plant Soil 384, 169–183.

Bajwa, A.A., Wang, H., Chauhan, B.S., Adkins, S.W., 2019. Effect of elevated carbon dioxide concentration on growth, productivity and glyphosate response of parthenium weed (Parthenium hysterophorus L.). Pest Manag. Sci. 75, 2934–2941.

Bannert, A., Kleineidam, K., Wissing, L., Mueller-Niggemann, C., Vogelsang, V., Welzl, G., Cao, Z., Schloter, M., 2011. Changes in diversity and functional gene abundances of microbial communities involved in nitrogen fixation, nitrification, and denitrification in a tidal wetland versus paddy soils cultivated for different time periods. Appl. Environ. Microbiol. 77. 10.1128/AEM.01751-10

Belnap, J., Phillips, S.L., Sherrod, S.K., Moldenke, A., 2005. Soil biota can change after exotic plant invasion: does this affect ecosystem processes? Ecology 86, 3007–3017.

Bengtson, P., Barker, J., Grayston, S.J., 2012. Evidence of a strong coupling between root exudation, C and N availability, and stimulated SOM decomposition caused by rhizosphere priming effects. Ecol. Evol. 2, 1843–1852.

Bergmann, J., Weigelt, A., van Der Plas, F., Laughlin, D.C., Kuyper, T.W., Guerrero-Ramirez, N., Valverde-Barrantes, O.J., Bruelheide, H., Freschet, G.T., Iversen, C.M., 2020. The fungal collaboration gradient dominates the root economics space in plants. Sci. Adv. 6, eaba3756.

Bezabih Beyene, B., Li, J., Yuan, J., Dong, Y., Liu, D., Chen, Z., Kim, J., Kang, H., Freeman, C., Ding, W., 2022. Non-native plant invasion can accelerate global climate change by increasing wetland methane and terrestrial nitrous oxide emissions. Glob. Chang. Biol. 28, 5453–5468.

Blumenthal, D.M., Resco, V., Morgan, J.A., Williams, D.G., LeCain, D.R., Hardy, E.M., Pendall, E., Bladyka, E., 2013. Invasive forb benefits from water savings by native plants and carbon fertilization under elevated CO 2 and warming. New Phytol. 200, 1156–1165.

Borchardt, M.A., Boehm, A.B., Salit, M., Spencer, S.K., Wigginton, K.R., Noble, R.T., 2021. The Environmental Microbiology Minimum Information (EMMI) Guidelines: qPCR and dPCR Quality and Reporting for Environmental Microbiology. Environ. Sci. Technol. 55, 10210–10223. 10.1021/acs.est.1c01767

Bradley, B.A., Blumenthal, D.M., Wilcove, D.S., Ziska, L.H., 2010. Predicting plant invasions in an era of global change. Trends Ecol. Evol. 25, 310–318.

Bremner, J.M., 1965. Inorganic forms of nitrogen. Methods soil Anal. Part 2 Chem. Microbiol. Prop. 9, 1179–1237.

Broadbent, A., Stevens, C.J., Peltzer, D.A., Ostle, N.J., Orwin, K.H., 2018. Belowground competition drives invasive plant impact on native species regardless of nitrogen availability. Oecologia 186, 577–587.

Butler, E.E., Mohanbabu, N., Wang, Z., Qiao, X., Isbell, F., Reich, P.B., 2025. Elevated CO2 and enriched nitrogen proportionally decrease species richness most at small spatial scales in a grassland experiment. J. Ecol. 113, 2800–2812.

Caldwell, B.A., 2005. Enzyme activities as a component of soil biodiversity: a review. Pedobiologia (Jena). 49, 637–644.

Chen, X., Chen, J., Le Roux, J.J., van Kleunen, M., van Groenigen, K.J., Fang, L., Hu, D., Fan, T., Liu, Y., Su, L., 2025. Global Patterns and Drivers of Soil Extracellular Enzyme Activities in Response to Plant Invasion: A Meta-Analysis. Glob. Ecol. Biogeogr. 34, e70084.

Cheng, W., Parton, W.J., Gonzalez-Meler, M.A., Phillips, R., Asao, S., McNickle, G.G., Brzostek, E., Jastrow, J.D., 2014. Synthesis and modeling perspectives of rhizosphere priming. New Phytol. 201, 31–44.

Choi, W.-J., Matushima, M., Ro, H.-M., 2011. Sensitivity of soil CO2 emissions to fertilizer nitrogen species: urea, ammonium sulfate, potassium nitrate, and ammonium nitrate. J. Korean Soc. Appl. Biol. Chem. 54, 1004–1007.

Cui, Y., Bing, H., Moorhead, D.L., Delgado-Baquerizo, M., Ye, L., Yu, J., Zhang, S., Wang, X., Peng, S., Guo, X., 2022. Ecoenzymatic stoichiometry reveals widespread soil phosphorus limitation to microbial metabolism across Chinese forests. Commun. Earth Environ. 3, 184.

Dafni, A., Heller, D., 1982. Adventive flora of Israel—Phytogeographical, ecological and agricultural aspects. Plant Syst. Evol. 140, 1–18.

Davidson, A.M., Jennions, M., Nicotra, A.B., 2011. Do invasive species show higher phenotypic plasticity than native species and, if so, is it adaptive? A meta-analysis. Ecol. Lett. 14, 419–431.

Davis, M.A., Grime, J.P., Thompson, K., 2000. Fluctuating resources in plant communities: a general theory of invasibility. J. Ecol. 88, 528–534.

Dijkstra, F.A., Carrillo, Y., Pendall, E., Morgan, J.A., 2013. Rhizosphere priming: a nutrient perspective. Front. Microbiol. 4, 216.

Dong, J., Hunt, J., Delhaize, E., Zheng, S.J., Jin, C.W., Tang, C., 2021. Impacts of elevated CO2 on plant resistance to nutrient deficiency and toxic ions via root exudates: a review. Sci. Total Environ. 754, 142434.

Drenovsky, R.E., Grewell, B.J., D’antonio, C.M., Funk, J.L., James, J.J., Molinari, N., Parker, I.M., Richards, C.L., 2012. A functional trait perspective on plant invasion. Ann. Bot. 110, 141–153.

Dukes, J.S., Mooney, H.A., 1999. Does global change increase the success of biological invaders? Trends Ecol. Evol. 14, 135–139.

Ehrenfeld, J.G., 2010. Ecosystem consequences of biological invasions. Annu. Rev. Ecol. Evol. Syst. 41, 59–80.

Ehrenfeld, J.G., 2003. Effects of exotic plant invasions on soil nutrient cycling processes. Ecosystems 6, 503–523.

Ekenler, M., Tabatabai, M.A., 2004. β-Glucosaminidase activity as an index of nitrogen mineralization in soils. Commun. Soil Sci. Plant Anal. 35, 1081–1094.

Emmett, B.D., Buckley, D.H., Drinkwater, L.E., 2020. Plant growth rate and nitrogen uptake shape rhizosphere bacterial community composition and activity in an agricultural field. New Phytol. 225, 960–973.

Enebe, M.C., Babalola, O.O., 2021. Soil fertilization affects the abundance and distribution of carbon and nitrogen cycling genes in the maize rhizosphere. AMB Express 11. 10.1186/s13568-021-01182-z

Fahey, C., Flory, S.L., 2022. Soil microbes alter competition between native and invasive plants. J. Ecol. 110, 404–414.

Fanin, N., Mooshammer, M., Sauvadet, M., Meng, C., Alvarez, G., Bernard, L., Bertrand, I., Blagodatskaya, E., Bon, L., Fontaine, S., 2022. Soil enzymes in response to climate warming: Mechanisms and feedbacks. Funct. Ecol. 36, 1378–1395.

Funk, J.L., Standish, R.J., Stock, W.D., Valladares, F., 2016. Plant functional traits of dominant native and invasive species in mediterranean-climate ecosystems. Ecology 97, 75–83.

Gao, G., Li, H., Shi, Y., Yang, T., Gao, C., Fan, K., Zhang, Y., Zhu, Y., Delgado-Baquerizo, M., Zheng, H., 2022. Continental-scale plant invasions reshuffle the soil microbiome of blue carbon ecosystems. Glob. Chang. Biol. 28, 4423–4438.

German, D.P., Weintraub, M.N., Grandy, A.S., Lauber, C.L., Rinkes, Z.L., Allison, S.D., 2011. Optimization of hydrolytic and oxidative enzyme methods for ecosystem studies. Soil Biol. Biochem. 43, 1387–1397.

Guo, X., Hu, Y., Ma, J.-Y., Wang, H., Wang, K.-L., Wang, T., Jiang, S.-Y., Jiao, J.-B., Sun, Y.-K., Jiang, X.-L., 2023. Nitrogen deposition effects on invasive and native plant competition: implications for future invasions. Ecotoxicol. Environ. Saf. 259, 115029.

Hager, H.A., Ryan, G.D., Newman, J.A., 2020. Effects of elevated CO2 on competition between native and invasive grasses. Oecologia 192, 1099–1110.

Han, X., Beck, K., Bürgmann, H., Frey, B., Stierli, B., Frossard, A., 2023. Synthetic oligonucleotides as quantitative PCR standards for quantifying microbial genes. Front. Microbiol. 14, 1279041.

Hellmann, J.J., Byers, J.E., Bierwagen, B.G., Dukes, J.S., 2008. Five potential consequences of climate change for invasive species. Conserv. Biol. 22, 534–543.

Hoppe, H.-G., 2018. Use of fluorogenic model substrates for extracellular enzyme activity (EEA) measurement of bacteria, in: Handbook of Methods in Aquatic Microbial Ecology. CRC Press, pp. 423–431.

Huo, C., Luo, Y., Cheng, W., 2017. Rhizosphere priming effect: a meta-analysis. Soil Biol. Biochem. 111, 78–84.

Incerti, G., Cartenì, F., Cesarano, G., Sarker, T.C., Abd El-Gawad, A.M., D’Ascoli, R., Bonanomi, G., Giannino, F., 2018. Faster N release, but not C loss, from leaf litter of invasives compared to native species in Mediterranean ecosystems. Front. Plant Sci. 9, 534.

Jin, H., Oduor, A.M.O., Xiao, L., Zhang, S., Liu, Y., 2025. Nutrient enrichment and interspecific competition modulate growth performance of invasive plant species regardless of nematodes. J. Plant Ecol. 18, rtaf060.

Jo, I., Fridley, J.D., Frank, D.A., 2017. Invasive plants accelerate nitrogen cycling: evidence from experimental woody monocultures. J. Ecol. 105, 1105–1110.

Kalra, Y.P., Maynard, D.C., 1991. Methods manual for forest soil and plant analysis.

Karkanis, A., Tsoutsoura, G., Ntanovasili, E., Mavroviti, V., Ntatsi, G., 2022. Bristly oxtongue (Helminthotheca echioides (L.) Holub) responses to sowing date, fertilization scheme, and chitosan application. Agronomy 12, 3028.

Kaštovská, E., Edwards, K., Picek, T., Šantrůčková, H., 2015. A larger investment into exudation by competitive versus conservative plants is connected to more coupled plant–microbe N cycling. Biogeochemistry 122, 47–59.

Kaushik, R., Sharma, M., Ramana, C. V, Sasikala, C., Pandit, M.K., 2023. Contrasting plant growth performance of invasive polyploid and native diploid Prosopis is mediated by the soil bacterial community. Ecol. Process. 12, 12.

Kelley, A.M., Fay, P.A., Polley, H.W., Gill, R.A., Jackson, R.B., 2011. Atmospheric CO2 and soil extracellular enzyme activity: a meta-analysis and CO2 gradient experiment. Ecosphere 2, 1–20.

Kuzyakov, Y., Razavi, B.S., 2019. Rhizosphere size and shape: temporal dynamics and spatial stationarity. Soil Biol. Biochem. 135, 343–360.

Lambers, H., Martinoia, E., Renton, M., 2015. Plant adaptations to severely phosphorus-impoverished soils. Curr. Opin. Plant Biol. 25, 23–31.

Lee, M.R., Flory, S.L., Phillips, R.P., 2012. Positive feedbacks to growth of an invasive grass through alteration of nitrogen cycling. Oecologia 170, 457–465.

Leishman, M.R., Haslehurst, T., Ares, A., Baruch, Z., 2007. Leaf trait relationships of native and invasive plants: community-and global-scale comparisons. New Phytol. 176, 635–643.

Li, J., He, J.-Z., Liu, M., Yan, Z.-Q., Xu, X.-L., Kuzyakov, Y., 2024. Invasive plant competitivity is mediated by nitrogen use strategies and rhizosphere microbiome. Soil Biol. Biochem. 192, 109361.

Li, J., You, Y., Zhang, W., Wang, Y., Liang, Y., Huang, H., Ma, H., He, Q., Ming, A., Huang, X., 2025. Soil microbial diversity and network complexity promote phosphorus transformation – a case of long-term mixed plantations of *Eucalyptus* and a nitrogen-fixing tree species. Biogeosciences 22, 4221–4239. 10.5194/bg-22-4221-2025

Liu, Y., Oduor, A.M.O., Zhang, Z., Manea, A., Tooth, I.M., Leishman, M.R., Xu, X., Van Kleunen, M., 2017. Do invasive alien plants benefit more from global environmental change than native plants? Glob. Chang. Biol. 23, 3363–3370.

Lu, X., Siemann, E., He, M., Wei, H., Shao, X., Ding, J., 2016. Warming benefits a native species competing with an invasive congener in the presence of a biocontrol beetle. New Phytol. 211, 1371–1381.

Markham, J., Chanway, C.P., 1996. Measuring plant neighbor effects. Funct. Ecol. 10, 549.

Matzrafi, M., Lazar, T.W., Sibony, M., Rubin, B., 2015. Conyza species: distribution and evolution of multiple target-site herbicide resistances. Planta 242, 259–267. 10.1007/s00425-015-2306-4

Meier, I.C., Tückmantel, T., Heitkötter, J., Müller, K., Preusser, S., Wrobel, T.J., Kandeler, E., Marschner, B., Leuschner, C., 2020. Root exudation of mature beech forests across a nutrient availability gradient: the role of root morphology and fungal activity. New Phytol. 226, 583–594.

Mori, T., 2024. Is enzymatic stoichiometry a reliable indicator of microbial limitations in carbon, nitrogen, or phosphorus? Sci. Total Environ. 955, 176928.

Negesse, Z., Pan, K., Guadie, A., Justine, M.F., Azene, B., Pandey, B., Wu, X., Sun, X., Zhang, L., 2025. Plant invasions alter soil biota and microbial activities: a global meta-analysis. Plant Soil 1–20.

Phillips, R.P., Meier, I.C., Bernhardt, E.S., Grandy, A.S., Wickings, K., Finzi, A.C., 2012. Roots and fungi accelerate carbon and nitrogen cycling in forests exposed to elevated CO2. Ecol. Lett. 15, 1042–1049.

Pugnaire, F.I., Morillo, J.A., Peñuelas, J., Reich, P.B., Bardgett, R.D., Gaxiola, A., Wardle, D.A., Van Der Putten, W.H., 2019. Climate change effects on plant-soil feedbacks and consequences for biodiversity and functioning of terrestrial ecosystems. Sci. Adv. 5, eaaz1834.

Pyšek, P., Hulme, P.E., Simberloff, D., Bacher, S., Blackburn, T.M., Carlton, J.T., Dawson, W., Essl, F., Foxcroft, L.C., Genovesi, P., 2020. Scientists’ warning on invasive alien species. Biol. Rev. 95, 1511–1534.

Qiu, J., 2015. A global synthesis of the effects of biological invasions on greenhouse gas emissions. Glob. Ecol. Biogeogr. 24, 1351–1362.

Ren, G.-Q., Li, Q., Li, Y., Li, J., Adomako, M.O., Dai, Z.-C., Li, G.-L., Wan, L.-Y., Zhang, B., Zou, C.B., 2019. The enhancement of root biomass increases the competitiveness of an invasive plant against a co-occurring native plant under elevated nitrogen deposition. Flora 261, 151486.

Sardans, J., Bartrons, M., Margalef, O., Gargallo-Garriga, A., Janssens, I.A., Ciais, P., Obersteiner, M., Sigurdsson, B.D., Chen, H.Y.H., Peñuelas, J., 2017. Plant invasion is associated with higher plant–soil nutrient concentrations in nutrient-poor environments. Glob. Chang. Biol. 23, 1282–1291.

Sinsabaugh, R.L., Hill, B.H., Follstad Shah, J.J., 2009. Ecoenzymatic stoichiometry of microbial organic nutrient acquisition in soil and sediment. Nature 462, 795–798.

Sinsabaugh, R.L., Lauber, C.L., Weintraub, M.N., Ahmed, B., Allison, S.D., Crenshaw, C., Contosta, A.R., Cusack, D., Frey, S., Gallo, M.E., 2008. Stoichiometry of soil enzyme activity at global scale. Ecol. Lett. 11, 1252–1264.

Sinsabaugh, R.L., Manzoni, S., Moorhead, D.L., Richter, A., 2013. Carbon use efficiency of microbial communities: stoichiometry, methodology and modelling. Ecol. Lett. 16, 930–939.

Sobuj, N., Singh, K., Byun, C., 2024. Responses of invasive and native plant species to drought stress and elevated CO2 concentrations: a meta-analysis. NeoBiota 96, 381–401.

Sorte, C.J.B., Ibáñez, I., Blumenthal, D.M., Molinari, N.A., Miller, L.P., Grosholz, E.D., Diez, J.M., D’Antonio, C.M., Olden, J.D., Jones, S.J., 2013. Poised to prosper? A cross-system comparison of climate change effects on native and non-native species performance. Ecol. Lett. 16, 261–270.

Te Beest, M., Esler, K.J., Richardson, D.M., 2015. Linking functional traits to impacts of invasive plant species: a case study. Plant Ecol. 216, 293–305.

Torres, N., Herrera, I., Fajardo, L., Bustamante, R.O., 2021. Meta-analysis of the impact of plant invasions on soil microbial communities. BMC Ecol. Evol. 21, 1–8.

Van der Putten, W.H., Bardgett, R.D., Bever, J.D., Bezemer, T.M., Casper, B.B., Fukami, T., Kardol, P., Klironomos, J.N., Kulmatiski, A., Schweitzer, J.A., 2013. Plant–soil feedbacks: the past, the present and future challenges. J. Ecol. 101, 265–276.

Wei, H., Yan, W., Quan, G., Zhang, J., Liang, K., 2017. Soil microbial carbon utilization, enzyme activities and nutrient availability responses to Bidens pilosa and a non-invasive congener under different irradiances. Sci. Rep. 7, 11309.

Weidenhamer, J.D., Callaway, R.M., 2010. Direct and indirect effects of invasive plants on soil chemistry and ecosystem function. J. Chem. Ecol. 36, 59–69.

Weltzin, J.F., Belote, R.T., Sanders, N.J., 2003. Biological invaders in a greenhouse world: will elevated CO2 fuel plant invasions? Front. Ecol. Environ. 1, 146–153.

Wen, Z., White, P.J., Shen, J., Lambers, H., 2022. Linking root exudation to belowground economic traits for resource acquisition. New Phytol. 233, 1620–1635.

Wright, I.J., Reich, P.B., Westoby, M., Ackerly, D.D., Baruch, Z., Bongers, F., Cavender-Bares, J., Chapin, T., Cornelissen, J.H.C., Diemer, M., 2004. The worldwide leaf economics spectrum. Nature 428, 821–827.

Xu, M., He, Z., Deng, Y., Wu, L., Van Nostrand, J.D., Hobbie, S.E., Reich, P.B., Zhou, J., 2013. Elevated CO2 influences microbial carbon and nitrogen cycling. BMC Microbiol. 13, 124.

Yanuka-Golub, K., Korenblum, E., Aronson, E.L., Matzrafi, M., 2025. Linking microbial-mediated methane production in wetlands to invasive plants. Soil Biol. Biochem. 109944.

Yu, H., Le Roux, J.J., Jiang, Z., Sun, F., Peng, C., Li, W., 2021. Soil nitrogen dynamics and competition during plant invasion: insights from Mikania micrantha invasions in China. New Phytol. 229, 3440–3452.

Zhang, T., Song, B., Wang, L., Li, Y., Wang, Y., Yuan, M., 2024. Spartina alterniflora invasion reduces soil microbial diversity and weakens soil microbial inter-species relationships in coastal wetlands. Front. Microbiol. 15, 1422534.

Zhang, Z., Pan, M., Zhang, X., Liu, Y., 2022. Responses of invasive and native plants to different forms and availability of phosphorus. Am. J. Bot. 109, 1560–1567.

Zheng, H., Vesterdal, L., Schmidt, I.K., Rousk, J., 2022. Ecoenzymatic stoichiometry can reflect microbial resource limitation, substrate quality, or both in forest soils. Soil Biol. Biochem. 167, 108613.

Zhou, Y., Staver, A.C., 2019. Enhanced activity of soil nutrient-releasing enzymes after plant invasion: a meta-analysis. Ecology 100, e02830.

Ziska, L.H., Blumenthal, D.M., Franks, S.J., 2019. Understanding the nexus of rising CO2, climate change, and evolution in weed biology. Invasive Plant Sci. Manag. 12, 79–88.

