## Supplemental Table S1 for "Warming and resource enrichment decouple growth from enzymatic investment, shifting the competitive balance between native and invasive plants"

**Table S1**. Primers and qPCR assay characteristics used in this study.

**$CO_2$**

| Gene | Target Organism/ Function | Primer Sequence (5'–3') | Amplicon Size (bp) | Standard / gBlock Info | Thermal Cycling Conditions | Efficiency (%) | $R^2$ |
|---|---|---|---|---|---|---|---|
| 16S | Bacteria (Biomass)[2] | F: CCGTCAATTCMTTTGAGTTT<br><br>R: CAGCMGCCGCGGTAANWC | bp 388 | *Geobacter metallireducens*genomic DNA (DSM 7210) Standard solution (103ng/µL): diluted twice with DW. Standard curve: 0.0051-51.5 ng/µL | 95°C 5m; x 40 cycles (95°C 30s, 64°C 30s, 72°C 30s) | 103%<br><br>102.1 % | 0.99<br><br>0.989 |
| nirS | Denitrifiers (Function)[1] | ***F: cd3AF***<br><br>GTS AAC GTS AAG GAR ACS GG<br><br>***R: R3cd***<br><br>GAS TTC GGR TGS GTC TTG A | 425 bp | Synthetic gBlock (IDT) | 95°C 3m; x 40 cycles (95°C 45s, 58°C 45s, 72°C 45s) | 101.7 %<br><br>98.1 % | 0.983<br><br>0.995 |
| amoA | Bacterial ammonia oxidizers (Function)[3] | F:<br><br>GGG GHT TYT ACT GGT GGT<br><br>R:<br><br>CCC CTC KGS AAA GCC TTC TTC | 491 bp | Synthetic gBlock (IDT) | 95°C 2m; 40x (95°C 45s, 55°C 45s, 72°C 45s) | 95.8 %<br><br>91.6 % | 0.99<br><br>0.99 |

**Fertilization**

| Gene | Target Organism/ Function | Primer Sequence (5'–3') | Amplicon Size (bp) | Standard / gBlock Info | Thermal Cycling Conditions | Efficiency (%) | $R^2$ |
|---|---|---|---|---|---|---|---|
| **16S** | Bacteria (Biomass)[2] | F: CCGTCAATTCMTTTGA GTTT<br>R: CAGCMGCCGCGGTAAN WC | bp 388 | *Geobacter metallireducensgenomic* DNA (DSM 7210) Standard solution (103ng/µL): diluted twice with DW. Standard curve: 0.002 - 20.6 ng/µL | 95°C 5m; 40x (95°C 30s, 64°C 30s, 72°C 30s) | 103.9%<br>97.9% | 0.99<br>0.984 |
| **nirS** | Denitrifiers (Function)[1] | F: cd3AF<br>GTS AAC GTS AAG GAR ACS GG<br>R: R3cd<br>GAS TTC GGR TGS GTC TTG A | ~425 bp | Synthetic gBlock (IDT) | 95°C 3m; 40x (95°C 45s, 58°C 45s, 72°C 45s) | 104.8%<br>96.1 | 0.98<br>0.982 |
| amoA | Bacterial ammonia oxidizers (Function)[3] | F:<br>GGG GHT TYT ACT GGT GGT<br>R:<br>CCC CTC KGS AAA GCC TTC TTC | 491 bp | Synthetic gBlock (IDT) | 95°C 2m; 40x (95°C 45s, 55°C 45s, 72°C 45s) | 95% | 0.984 |

### Temperature

| Gene | Target Organism/ Function | Primer Sequence (5'–3') | Amplicon Size (bp) | Standard / gBlock Info | Thermal Cycling Conditions | Efficiency (%) | $R^2$ |
|---|---|---|---|---|---|---|---|
| **16S** | Bacteria (Biomass)[2] | F: CCGTCAATTCMTTT GAGTTT<br>R: CAGCMGCCGCGGTA ANWC | bp 388 | *Geobacter metallireducens genomic* DNA (DSM 7210) Standard solution (103ng/µL): diluted twice with DW. Standard curve*:* 0.002 - 20.6 ng/µL | 95°C 5m; 40x (95°C 30s, 64°C 30s, 72°C 30s) | 103.9% | 0.99 |
| **nirS** | Denitrifiers (Function)[1] | F: cd3AF<br>GTS AAC GTS AAG GAR ACS GG<br>R: R3cd<br>GAS TTC GGR TGS GTC TTG A | ~425 bp | Synthetic gBlock (IDT) | 95°C 3m; 40x (95°C 45s, 58°C 45s, 72°C 45s) | 95.6% | 0.984 |
| amoA | Bacterial ammonia oxidizers (Function)[3] | F: GGG GHT TYT ACT GGT GGT<br>R: CCC CTC KGS AAA GCC TTC TTC | 491 bp | Synthetic gBlock (IDT) | 95°C 2m; 40x (95°C 45s, 55°C 45s, 72°C 45s) | 93.8% | 0.987 |

1. Throbäck, I. N., Enwall, K., Jarvis, Å. & Hallin, S. Reassessing PCR primers targeting nirS, nirK and nosZ genes for community surveys of denitrifying bacteria with DGGE. *FEMS Microbiol. Ecol.* 49, 401–417 (2004).

2 Iasur-Kruh LHadar Y, Minz D2011.Isolation and Bioaugmentation of an Estradiol-Degrading Bacterium and Its Integration into a Mature Biofilm. Appl Environ Microbiol77:.https://doi.org/10.1128/AEM.00691-11

3 Rotthauwe, J. H., Witzel, K. P., & Liesack, W. The ammonia monooxygenase structural gene amoA as a functional marker: molecular fine-scale analysis of natural ammonia-oxidizing populations. Appl. Environ. Microbiol. 63, 4704–4712 (1997)

***Table S1 Notes***: Quantitative PCR (qPCR) was performed using the SsoAdvanced™ Universal Inhibitor-Tolerant SYBR® Green Supermix (Bio-Rad) on a Bio-Rad CFX96 Real-Time PCR System. Gene Target represents the specific genetic marker for taxonomic (16S) or functional (nirS) potential. Amplicon denotes the expected size of the PCR product in base pairs (bp). Standard Source refers to the material used for the 10-fold serial dilutions used to generate standard curves; nirS standards were generated using synthetic double-stranded DNA fragments (gBlocks, Integrated DNA Technologies). Thermal Cycling includes the initial denaturation followed by the number of cycles and specific temperature/time for denaturation, annealing, and extension. Efficiency was calculated from the slope of the standard curve according to the formula $E = 10^{(-1/slope)} - 1$. All samples, standards, and non-template controls (NTC) were run in triplicate.
