## Supplemental Table S2 for "Warming and resource enrichment decouple growth from enzymatic investment, shifting the competitive balance between native and invasive plants"

**Table S2.** Standard curve and assay parameters for extracellular enzyme activities.

$CO_2$

| Enzyme | Substrate | Standard-MUF concentration range (µM) | Incubation time (h) | Standard Curve $R^2$ | Mean quench coefficient (%) |
|---|---|---|---|---|---|
| α-1,4-glucosidase (AG; EC 3.2.1.20) | 4-MUF-α-D-glucopyranoside (69591) | 0–681 | 1 | 0.977±0.0153 | * 0.577±0.127 |
| β-1,4-N-acetylglucosaminidase (NAGase; EC 3.2.1.52) | 4-MUF- N-acetyl-β-D-glucosaminide (M2133) | 0–681 | 1 | 0.977±0.0153 | *0.577±0.127 |

Fertilizer

| Enzyme | Substrate | Standard-MUF concentration range (µM) | Incubation time (h) | Standard Curve $R^2$ | Mean quench coefficient (%) |
|---|---|---|---|---|---|
| α-1,4-glucosidase (AG; EC 3.2.1.20) | 4-MUF-α-D-glucopyranoside (69591) | 0–681 | 1 | 0.977±0.0143 | * 0.513±0.158 |
| β-1,4-N-acetylglucosaminidase (NAGase; EC 3.2.1.52) | 4-MUF- N-acetyl-β-D-glucosaminide (M2133) | 0–681 | 1 | 0.977±0.0143 | *0.513±0.158 |

Temperature

| Enzyme | Substrate | Standard-MUF concentration range (µM) | Incubation time (h) | Standard Curve $R^2$ | Mean quench coefficient (%) |
|---|---|---|---|---|---|
| α-1,4-glucosidase (AG; EC 3.2.1.20) | 4-MUF-α-D-glucopyranoside (69591) | 0–681 | 1 | 0.977±0.0130 | * 0.597±0.216 |
| β-1,4-N-acetylglucosaminidase (NAGase; EC 3.2.1.52) | 4-MUF- N-acetyl-β-D-glucosaminide (M2133) | 0–681 | 1 | 0.977±0.0130 | *0.597±0.216 |

*Note: Quench coefficients represent the ratio of fluorescence in the soil homogenate to fluorescence in sterile water, used to correct for soil-specific signal attenuation.

*Calculations were based on the molar mass of neutral MUF (176.17 g/mol) to ensure accurate stoichiometric reporting of product release.
